# A genetic variant of the Wnt receptor LRP6 accelerates synapse degeneration during ageing and in Alzheimer’s disease

**DOI:** 10.1101/2022.04.06.487208

**Authors:** Megan E. Jones, Johanna Büchler, Tom Dufor, Katharina Boroviak, Emmanouil Metzakopian, Alasdair Gibb, Patricia C. Salinas

**Author notes:** Contributed equally to this work (Megan E. Jones, Johanna Büchler). Corresponding author, Correspondence: Professor Patricia C. Salinas.

## Abstract

Synapse loss strongly correlates with cognitive decline in Alzheimer’s Disease (AD), but the underlying mechanisms are poorly understood. Studies suggest that deficient Wnt signalling, a pathway required for neuronal connectivity, contributes to synapse dysfunction and loss in AD. Consistent with this idea, a variant of *Lrp6, (Lrp6-val)*, which confers reduced Wnt signalling, has been linked to late onset AD. However, the impact of *Lrp6-val* on synapses in the healthy and AD brain has not been examined. Using CRISPR/Cas9 genome editing, we generated a novel knock-in mouse model carrying this *Lrp6* variant to study its role in synaptic integrity. *Lrp6-val* mice develop normally and do not exhibit morphological brain abnormalities. Hippocampal neurons from *Lrp6-val* mice do not respond to Wnt7a, a Wnt ligand that promotes synaptic assembly through the Frizzled-5 (Fz5) receptor. Activation of the Wnt pathway by Wnt ligands leads to the formation of a complex between LRP6 and Fz5. In contrast, LRP6-Val impairs the formation of the LRP6-Fz5 complex elicited by Wnt7a, as detected by proximity ligation assay (PLA). We demonstrate that *Lrp6-val* mice exhibit structural and functional synaptic defects that become more pronounced with age, consistent with decreased canonical Wnt signalling during ageing. To investigate the contribution of this variant to AD, *Lrp6-val* mice were crossed to *hAPP^NL-G-F/NL-G-F^ (NL-G-F*), a knock-in AD mouse model. The presence of the *Lrp6-val* variant significantly exacerbates synapse loss around amyloid-β plaques in *NL-G-F* mice. Our findings uncover a novel role for the *Lrp6-val* variant in synapse vulnerability during ageing and its contribution to synapse degeneration in AD.

## Introduction

In Alzheimer’s Disease (AD), memory impairment strongly correlates with synapse degeneration (Terry et al. 1991; Scheff & Price 2006; Scheff et al. 2007; Shankar & Walsh 2009). Synaptic changes occur early in the disease, before amyloid-β (Aβ) plaque formation and neuronal loss (Shankar & Walsh 2009; Mucke & Selkoe 2012; Marsh & Alifragis 2018). At later stages in the disease, further synapse loss is observed around Aβ plaques (Dong et al. 2007; Koffie et al. 2009; Koffie et al. 2012; Jackson et al. 2016). It is well documented that accumulation of oligomeric forms of Aβ triggers synapse dysfunction and degeneration (Shankar & Walsh 2009; Mucke & Selkoe 2012; Selkoe & Hardy 2016; Marsh & Alifragis 2018). Numerous studies demonstrate that Aβ oligomers interfere with signalling pathways, which are critical for maintaining synapse integrity, resulting in synapse weakening and loss (Lin & Koleske 2010; Chen et al. 2019). However, the molecular mechanisms that lead to synapse dysfunction and loss in AD remain poorly understood.

The canonical Wnt signalling pathway, required for synapse function and stability, is impaired in AD (**Figure 1A**) (Purro et al. 2012; Liu et al. 2014; Marzo et al. 2016). Studies on Wnt ligands, Frizzled receptors, and the co-receptor LRP6 demonstrate their critical role in synaptogenesis (Ciani et al. 2011; Sharma et al. 2013; McLeod et al. 2018), and in synapse integrity in the adult brain (Galli et al. 2014; Liu et al. 2014; Marzo et al. 2016). The first piece of evidence that Wnt signalling is compromised in AD came from the finding that Dickkopf-1 (Dkk1), a secreted Wnt antagonist, is elevated in the brains of AD patients and in AD mouse models (Caricasole et al. 2004; Rosi et al. 2010). Notably, Dkk1 is required for Aβ-induced synapse degeneration as blockade of Dkk1 protects against Aβ-mediated synapse loss (Purro et al. 2012; Sellers et al. 2018). Consistently, *in vivo* expression of Dkk1 in the adult brain induces synapse loss, LTP defects (Marzo et al. 2016) and memory impairment (Killick et al. 2014; Marzo et al. 2016), as observed in AD mouse models. Second, conditional knockout (cKO) of *Lrp6* in an AD mouse model increases amyloid pathology and exacerbates cognitive deficits (Liu et al. 2014). Third, a genetic link between deficient Wnt signalling and AD came from the identification of three genetic variants of *Lrp6* associated with late onset AD (LOAD) (De Ferrari et al. 2007; Alarcón et al. 2013). Notably, a nonsynonymous SNP (rs2302685) (**Figure 1B**), which has an allele frequency of 0.17 in the European population (1000 Genomes Project Consortium et al. 2015), results in a conservative substitution of isoleucine to valine at amino acid 1062 (*Lrp6^Ile-1062-Val^; Lrp6-val herein*). This substitution is located in the fourth β-propeller of the extracellular domain where some Wnt ligands bind (Bourhis et al. 2010; Chen et al. 2011; Hirai et al. 2019) (**Figure 1B**). The *Lrp6-val* variant confers reduced Wnt signalling in cell lines in response to a Wnt ligand (De Ferrari et al. 2007). However, the impact of this variant on brain development, neuronal connectivity and amyloid pathology remains unexplored.

**Figure 1.**
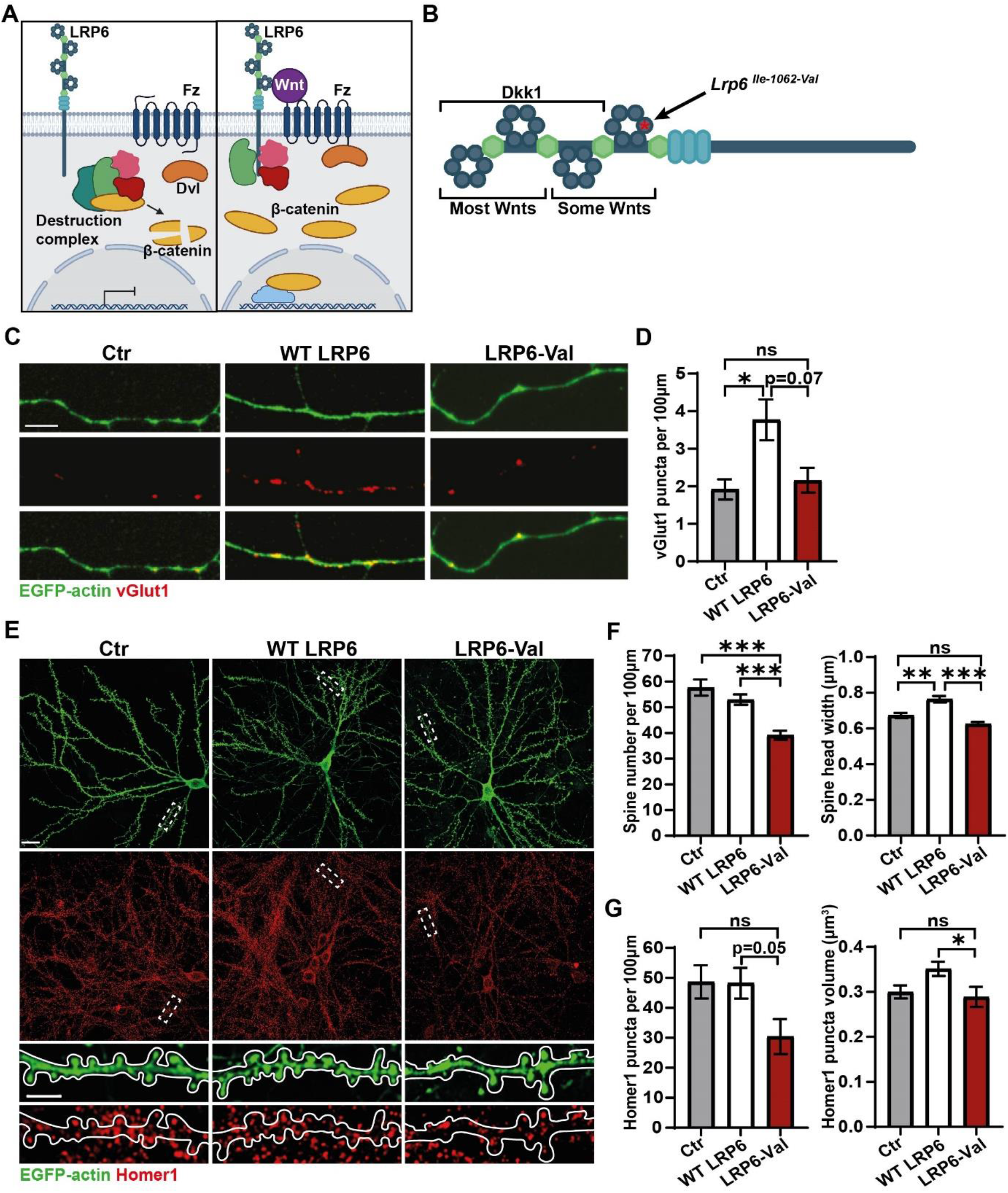
LRP6-Val gain-of-function induces synaptic defects in cultured neurons. A) Diagram of canonical Wnt signalling showing the role of LRP6 in this pathway. The left panel shows that in the absence of Wnt, the LRP6 co-receptor does not form a complex with Frizzled receptors. β-catenin is sequestered and degraded by the destruction complex preventing transcription of Wnt target genes. The right panel shows that Wnt, LRP6 and Fz receptors form a complex required for downstream signalling. Activation of the pathway results in Dishevelled (Dvl) recruitment to the plasma membrane and disassembly of the destruction complex. β-catenin accumulates and translocates to the nucleus enabling transcription of Wnt target genes. B) Schematic representation of the LRP6 protein showing the location of the *Lrp6-val* SNP (red asterisk and arrow) and the areas where the Wnt antagonist Dkk1 and Wnt ligands bind to. C) Confocal images of vGlut1 (red) puncta on isolated axons of neurons expressing EGFP-actin alone or EGFP-actin and human WT LRP6 or human LRP6-Val. Scale bar = 5 μm. D) Quantification of vGlut1 puncta density showed that WT LRP6 promoted the assembly of presynaptic sites but LRP6-Val did not. N = 4 independent cultures, 10-12 axons imaged per culture. Kruskal Wallace with Dunn’s post-hoc test. * p < 0.05. E) Top: Confocal images of Homer1 (red) and GFP (green) of neurons expressing EGFP-actin and human WT LRP6 or human LRP6-Val. Scale bar = 21 μm. Bottom: Higher magnification of areas of interest show dendritic spines (GFP; green) and Homer1 (red) puncta along dendrites. Scale bar = 5 μm. F) Left panel: Gain-of-function of LRP6-Val resulted in reduced spine density. 3 independent cultures, 8-10 cells imaged per culture. One-way-ANOVA with Tukey’s post-hoc test. *** p < 0.001. Right panel: Expression of LRP6-Val failed to increase spine size. 3 independent cultures, 8-10 cells imaged per culture. Kruskal-Wallis with Dunn’s post-hoc. ** p < 0.01; *** p < 0.001. G) LRP6-Val expression led to smaller Homer1 puncta. 2 independent cultures, 8-10 cells imaged per culture. One-way-ANOVA with Tukey’s post-hoc test. * p < 0.05. Data are represented as mean ± SEM.

Here, we investigated the impact of the *Lrp6-val* variant on the adult and ageing hippocampus and in the pathogenesis of AD by generating a novel knock-in mouse model using CRISPR/Cas9 genome editing. Homozygous *Lrp6-val* mice developed normally but showed structural and functional synaptic defects in the hippocampus that became more pronounced with age. In these mice, we observed decreased levels of canonical Wnt signalling with age. Notably, neurons from *Lrp6-val* mice were unable to respond to Wnt7a to promote synapse formation. As Wnt ligands promote the interaction between LRP6 and Frizzled receptors (Angers & Moon 2009; MacDonald et al. 2009; Hua et al. 2018), we examined if the LRP6-Val variant affects this interaction by focusing on Frizzled5 (Fz5), a Wnt7a receptor required for presynaptic assembly in hippocampal neurons (Sahores et al. 2010). We found that Wnt7a increased the association between wildtype LRP6 and Fz5, whereas this interaction was significantly impaired in the presence of LRP6-Val. Next, we examined the contribution of the *Lrp6-val* variant to AD pathogenesis by crossing these mice to *hAPP^NL-G-F/NL-G-F^ (NL-G-F*), a knock-in AD mouse model. The *Lrp6-val* variant significantly increased synapse degeneration in *NL-G-F* mice. Together, our findings uncover a novel function for the *Lrp6-val* variant in synapse degeneration during ageing and in AD. Our results demonstrate, for the first time, that the valine substitution in the extracellular domain of LRP6 impairs its interaction with Fz5, mediated by Wnt7a, and therefore affects downstream signalling.

## Results

### LRP6-Val fails to stimulate synaptic assembly and induces spine loss

LRP6 is required for synapse formation and synapse maintenance (Sharma et al. 2013; Liu et al. 2014). Furthermore, the *Lrp6-val* variant is linked to late onset AD (LOAD) (De Ferrari et al. 2007), but its impact on neuronal connectivity has not been explored. To assess the effect of LRP6-Val at synapses, we expressed human wildtype LRP6 (WT LRP6) or human LRP6-Val in cultured hippocampal neurons. Expression of WT LRP6 increased the number of vGlut1 puncta along axons (**Figures 1C and 1D**). In contrast, LRP6-Val failed to increase the number of presynaptic sites above control cells (**Figures 1C and 1D**). Thus, the LRP6-Val variant is unable to induce presynaptic assembly in neurons.

Next, we examined the impact of LRP6-Val on dendritic spines (**Figure 1E**). Expression of WT LRP6 increased spine head width but had no effect on spine density (**Figures 1E and 1F**). In contrast, expression of LRP6-Val failed to increase spine head width and decreased spine density when compared to control and WT LRP6 expressing cells (**Figures 1E and 1F**). Thus, gain-of-function of LRP6-Val is unable to promote dendritic spine growth, whilst inducing spine loss. A strong trend (p=0.05) towards fewer Homer1 puncta was observed in LRP6-Val expressing neurons compared to WT LRP6 expressing or control neurons (**Figures 1E and 1G**).

Although no differences in Homer1 puncta volume were found between control and WT LRP6 cells (**Figures 1E and 1G**), a significant decrease was observed in LRP6-Val expressing neurons compared to those expressing WT LRP6 (**Figures 1E and 1G**). Thus, gain-of-function of LRP6-Val in neurons decreases the formation of both pre- and post-synaptic sites compared to WT LRP6.

### *Lrp6-val* knock-in mice develop normally and LRP6-Val does not affect its synaptic localisation

To investigate the *in vivo* role of the *Lrp6-val* variant, we generated a novel knock-in mouse model using CRISPR/Cas9 genome editing. The *Lrp6* A→G point mutation, which results in the substitution of isoleucine to valine, was introduced at the endogenous *Lrp6* locus in the mouse genome, creating a mouse line that carries the *Lrp6-val* variant globally. DNA sequencing confirmed the successful generation of both heterozygous (*Lrp6-val* het) and homozygous (*Lrp6-val* hom) knock-in animals (**Figure 2A**). *Lrp6-val* het and *Lrp6-val* hom mice developed normally with no visible morphological abnormalities and no differences in weight (**Figures S1A S1B,** and data not shown). Furthermore, *Lrp6-val* het and *Lrp6-val* hom mice exhibit a similar synaptic phenotype (see below). Herein, we mainly examined *Lrp6-val* hom (*Lrp6-val*) mice.

**Figure 2.**
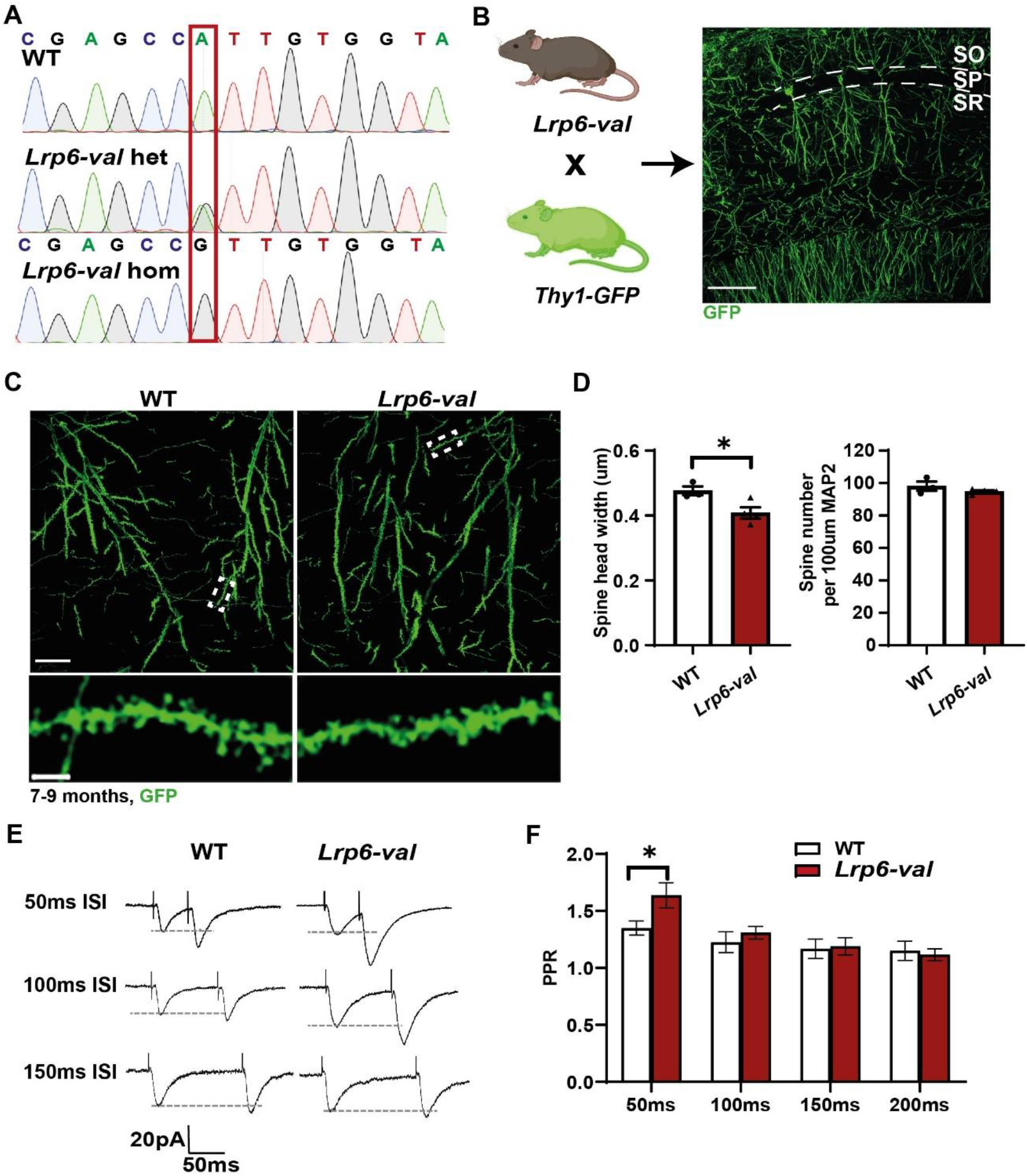
*Lrp6-val* mice exhibit synaptic defects at 7-9 months. A) Sanger trace examples of WT, *Lrp6-val* heterozygous (*Lrp6-val* het) and *Lrp6-val* homozygous (*Lrp6-val* hom) knock-in mice. B) Dendritic spines were analysed in *Lrp6-val* hom knock-in mice crossed to a *Thy1-GFP*line. SO = stratum oriens, SP = stratum pyramidale, SR = stratum radiatum. Scale bar = 100 μm. C) Confocal images of apical dendrites of CA1 pyramidal neurons of WT and *Lrp6-val* mice at 7-9 months. Scale bar = 25 μm. Inset shows spines along a dendrite. Scale bar = 3 μm. D) *Lrp6-val* mice display reduced spine head width. WT N = 3 *Lrp6-val* N = 4. Unpaired T-test. * p < 0.05. E) Representative paired-pulse recordings of synaptic currents from WT and *Lrp6-val* brain slices at different inter-stimulus intervals (ISIs). F) Graph displays the mean PPR from all recorded cells. *Lrp6-val* increased the PPR a 50 ms inter-stimulus intervals. N *=* 11-21 cells recorded from 4-5 animals per genotype. Repeated measures one-way-ANOVA with Tukey’s post-hoc test. * p < 0.05. Data are represented as mean ± SEM.

We next investigated whether the *Lrp6-val* variant was differentially expressed or affected its synaptic localisation. The mRNA and protein levels of LRP6 in the hippocampus of *Lrp6-val*mice were the same (**Figures S1C-S1E**). For its synaptic localisation, we used structured-illumination microscopy (SIM). Both WT LRP6 and LRP6-Val were present at approximately 80% of excitatory synapses (**Figures S1F and S1G**) but no differences in LRP6 localisation were observed between neurons from WT and *Lrp6-val* mice (**Figure S1G**). Thus, carrying the *Lrp6-val* variant does not alter the synaptic localisation of this receptor.

### *Lrp6-val* causes progressive synaptic defects with age

Spine formation and spine growth are modulated by Wnt signalling in the hippocampus (Ciani et al. 2011; Chen et al. 2017; McLeod et al. 2018; Ramos-Fernández et al. 2019). Given that expression of LRP6-Val failed to stimulate spine growth and induced spine loss in cultured neurons (**Figures 1E and 1F**), we examined the *in vivo* impact of *Lrp6-val* on these post-synaptic structures. We analysed dendritic spines on the apical dendrites of CA1 pyramidal neurons in adult *Lrp6-val* knock-in mice crossed to a *Thy1-GFP* expressing line (Feng et al. 2000) (**Figure 2B**). Although no differences in spine density were observed, spine head width was reduced in *Lrp6-val* mice compared to WT mice at 7-9 months (**Figures 2C and 2D**). Thus, adult mice carrying the *Lrp6-val* variant display impaired spine growth.

Given the synaptic localisation of LRP6 and the role of Wnt signalling in synaptic function (Dickins & Salinas 2013; McLeod & Salinas 2018; Oliva et al. 2018), we examined synaptic transmission in the *Lrp6-val* mice at Schaffer collateral (SC)-CA1 synapses, where deficient Wnt signalling leads to defects in synaptic transmission (Marzo et al. 2016). Evoked excitatory post-synaptic currents (EPSCs), in response to SC stimulation of increasing intensity (inputoutput (I/O) curve), were recorded in CA1 pyramidal cells at 7-9 months. No significant differences were observed between WT and *Lrp6-val* mice even at high stimulation intensities (**Figure S2A and S2B**), suggesting that basal synaptic transmission at SC-CA1 synapses is unaffected by the presence of the *Lrp6-val* variant at this age.

Wnt signalling deficient mice exhibit reduced neurotransmitter release probability (Galli et al. 2014; Ciani et al. 2015). We therefore investigated possible defects in neurotransmitter release. EPSCs evoked at brief intervals at SC-CA1 synapses were recorded from WT and *Lrp6-val* mice at 7-9 months. We analysed the paired pulse ratio (PPR), which depends on presynaptic short-term plasticity mechanisms and is inversely correlated with release probability (Dobrunz & Stevens 1997; Fioravante & Regehr 2011). *Lrp6-val* mice exhibited increased PPRs compared to WT mice at 50 ms inter-stimulus intervals (ISI), consistent with a reduced release probability at 7-9 months (**Figures 2E and 2F**). Thus, neurotransmitter release is compromised in adult *Lrp6-val* mice.

Given that cKO mice for *Lrp6* exhibit age-dependent synaptic deficits (Liu et al. 2014), we interrogated whether synaptic defects become more pronounced with age. We evaluated dendritic spines in the *Lrp6-val;Thy1-GFP* mice with age (**Figure 3A**). Our analyses revealed a significant decrease in both spine density and head width in *Lrp6-val* mice compared to WT mice at 12-14 months (**Figure 3B**). Given that spine size but not spine number was affected in 7-9-month-old *Lrp6-val* mice, these results demonstrate that spine deficits become more severe with age.

**Figure 3.**
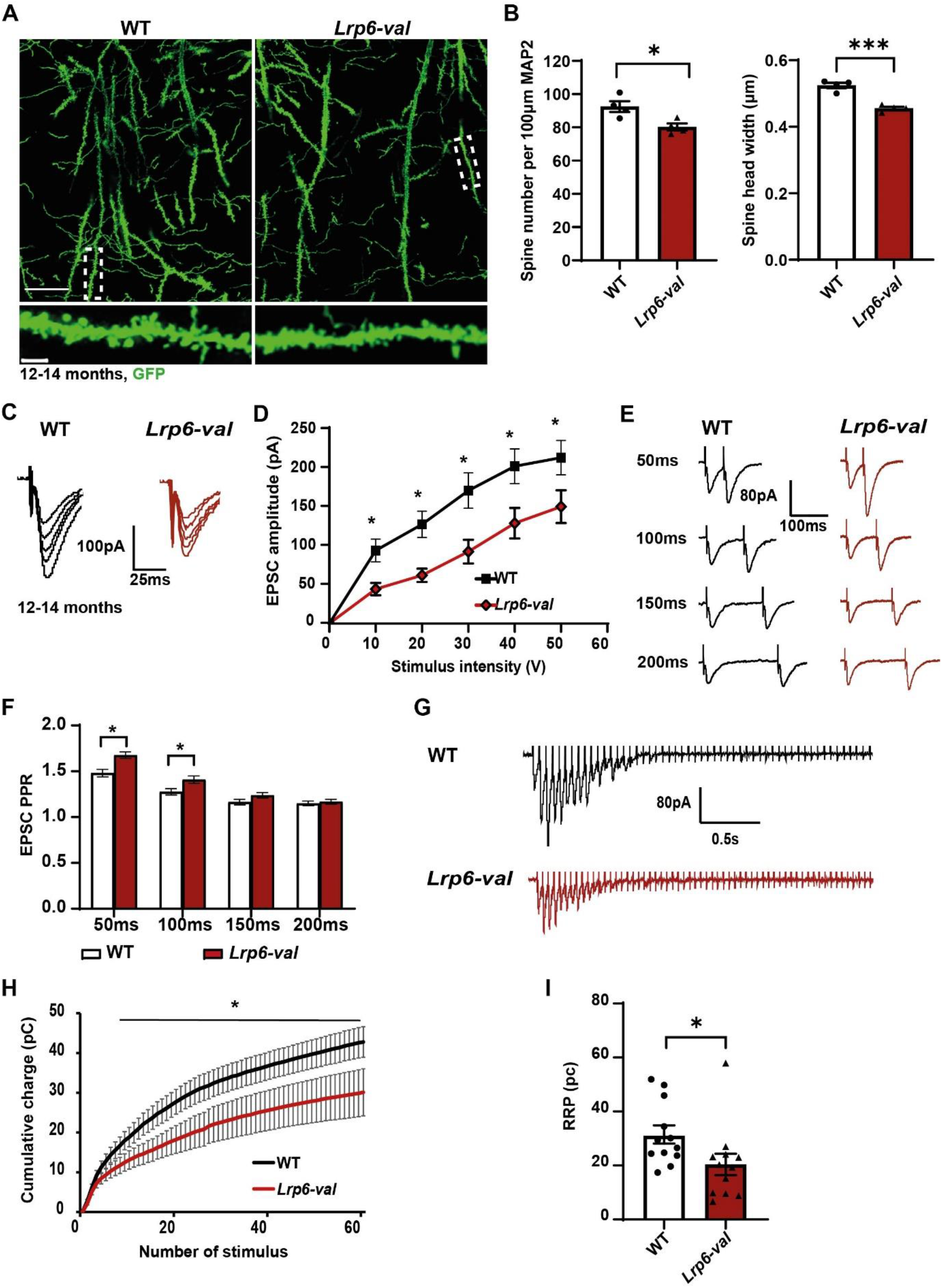
*Lrp6-val* mice display postsynaptic defects and impaired basal synaptic transmission, synaptic vesicle release and RRP size at 12 months. A) Top: Dendritic spines were analysed at 12-14 months in *Lrp6-val* knock-in mice crossed to a *Thy1-GFP* line. Scale bar = 25 μm. Bottom: Confocal images of regions of interest containing apical dendrites of CA1 pyramidal neurons in WT and *Lrp6-val* mice. Scale bar = 3 μm. B) *Lrp6-val* mice have smaller and fewer spines. WT N = 4, *Lrp6-val* N = 4. Unpaired T-test. * p < 0.05, *** p < 0.001. C) Representative traces of evoked excitatory post-synaptic currents (EPSCs) elicited at increasing stimulation voltages with an average of three responses for each stimulus voltage (10 V; 20 V; 30 V; 40 V and 50 V). D) Input-output curves showing a significant reduction in EPSC amplitude in hippocampal slices from *Lrp6-val* mice. N = 12-13 cells recorded from 4 animals per genotype. Repeated measures one-way-ANOVA with Tukey’s post-hoc test. * p < 0.05. E) Representative traces of paired pulse evoked EPSCs at different interstimulus intervals using brain slices from WT and *Lrp6-val* mice. F) Graph displays the mean PPR from all cells. *Lrp6-val* mice display increased PPR at 50 ms and 100 ms ISI. N = 13-14 cells from 4 animals per genotype. Repeated measure one-way-ANOVA with Tukey’s post-hoc test. * p < 0.05. G) Representative traces of EPSCs elicited by a 20 Hz electrical stimulation for 3 s recorded from WT and *Lrp6-val* mice. H) Graph showing reduced mean cumulative charge in *Lrp6-val* mice. N = 12 cells from 4 animals per genotype. Repeated measure one-way-ANOVA with Tukey’s post-hoc test. * p < 0.05. I) Graph displays the readily releasable pool (RRP) size, obtained from all cells. *Lrp6-val* mice exhibited a reduced RRP. N = 12 cells from 4 animals per genotype. Unpaired Student’s T-test. * p < 0.05. Data are represented as mean ± SEM.

Next, we evaluated basal synaptic transmission in 12-month-old mice by performing I/O curve recordings in hippocampal slices. Interestingly, the amplitude of evoked EPSCs were significantly smaller in 12-month-old *Lrp6-val* mice compared to WT mice at all stimulus intensities (**Figures 3C and 3D**). Thus, the *Lrp6-val* variant impairs basal synaptic transmission at this age but not at 7-9 months (**Figure S2A and S2B**). We then measured neurotransmitter release probability and found that the PPR was significantly higher in *Lrp6-val* mice compared to WT mice at 50 and 100 ms ISI, consistent with a reduction in release probability (**Figures 3E and 3F**). Given that PPR was significantly higher at 50 ms but not at 100 ms ISI in *Lrp6-val* mice at 7-9 months, these results suggest that defects in neurotransmitter release are more pronounced in older animals.

To further define the role of *Lrp6-val* in neurotransmitter release at 12 months, we recorded the responses to a 3 second high-frequency stimulus train (20Hz), which fully depletes presynaptic terminals of the readily releasable pool (RRP) (Wesseling & Lo 2002; Stevens & Williams 2007). Using first order correction for vesicle recycling (Wesseling & Lo 2002), a significant reduction in the size of the RRP was observed in 12-month-old *Lrp6-val* mice when compared to WT mice (**Figures 3G, 3H and 3I**). However, synaptic vesicle fusion efficiency and recycling rate were not affected (**Figure S2C**), suggesting that the defect in release probability is due to a reduced RRP.

The defects in vesicular release probability at 12 months suggested possible structural changes at presynaptic terminals of *Lrp6-val* mice. To investigate this, we evaluated the number and size of presynaptic boutons in *Lrp6-val* mice at 12-14 months in the CA1 stratum radiatum (SR) using the presynaptic marker, vGlut1 (**Figure 4A**). Indeed, *Lrp6-val* mice had smaller and fewer vGlut1 puncta compared to WT mice (**Figure 4B**). As *Lrp6-val* mice exhibited deficits in synaptic vesicle release, due to a smaller RRP, we assessed possible ultrastructural changes by electron microscopy (EM). We observed fewer synaptic vesicles at presynaptic terminals, but no differences in the length of the post-synaptic density, of *Lrp6-val* mice at SC-CA1 synapses (**Figures 4C and 4D**). These findings indicate that impaired neurotransmitter release in *Lrp6-val* mice is due to a reduction in synaptic vesicle number, consistent with a reduced RRP.

**Figure 4.**
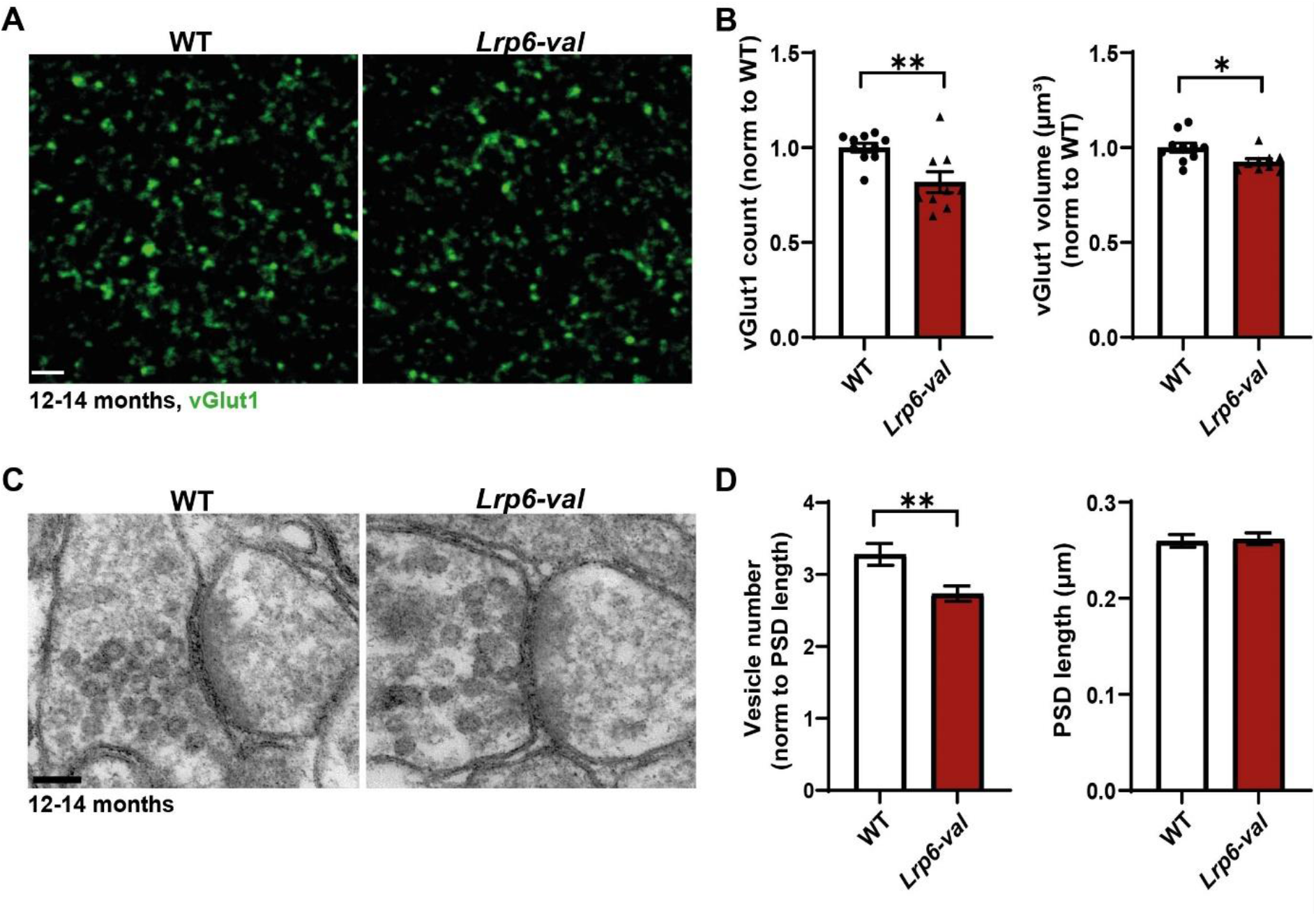
Presynaptic defects of *Lrp6-val* mice at 12-14 months. A) Confocal images of vGlut1-labelled excitatory presynaptic terminals in the CA1 stratum radiatum area of WT and *Lrp6-val* mice at 12-14 months. Scale bar = 2 μm. B) *Lrp6-val* mice had fewer and smaller vGlut1 puncta. WT N = 10 *Lrp6-val* N = 9. Unpaired T-test. * p < 0.05, ** p < 0.01. C) Electron microscopy images of an excitatory synapse of 12-14-month-old WT and *Lrp6-val* mice. Scale bar = 100 nm. D) *Lrp6-val* mice had fewer synaptic vesicles but no changes in PSD length were observed. N = 5, 19-25 images per animal. Mann-Whitney test. ** p < 0.01. Data are represented as mean ± SEM.

Given that defects at the pre- and post-synaptic sites were exacerbated with age, we compared the number of excitatory synapses at the different ages. Synapses were quantified based on the co-localisation of Bassoon and Homer1 in the CA1 SR region. At 7-9 months and 12 months, no differences were observed between WT and *Lrp6-val* mice (**Figures 5A–5D**). However, a significant reduction in the number of excitatory synapses was detected at 16-18 months in *Lrp6-val* mice when compared to WT mice (**Figures 5E and 5F**). Due to the age of these animals, we examined possible neuronal loss, which could affect synapse number, in the stratum pyramidale layer using DAPI and NeuN (**Figure S3A**). No differences in the percentage of NeuN positive cells were identified between WT and *Lrp6-val* mice at 16-18 months (**Figure S3B**). Thus, the *Lrp6-val* variant confers increased synaptic vulnerability as animals age, a process that is not due to changes in neuronal number.

**Figure 5.**
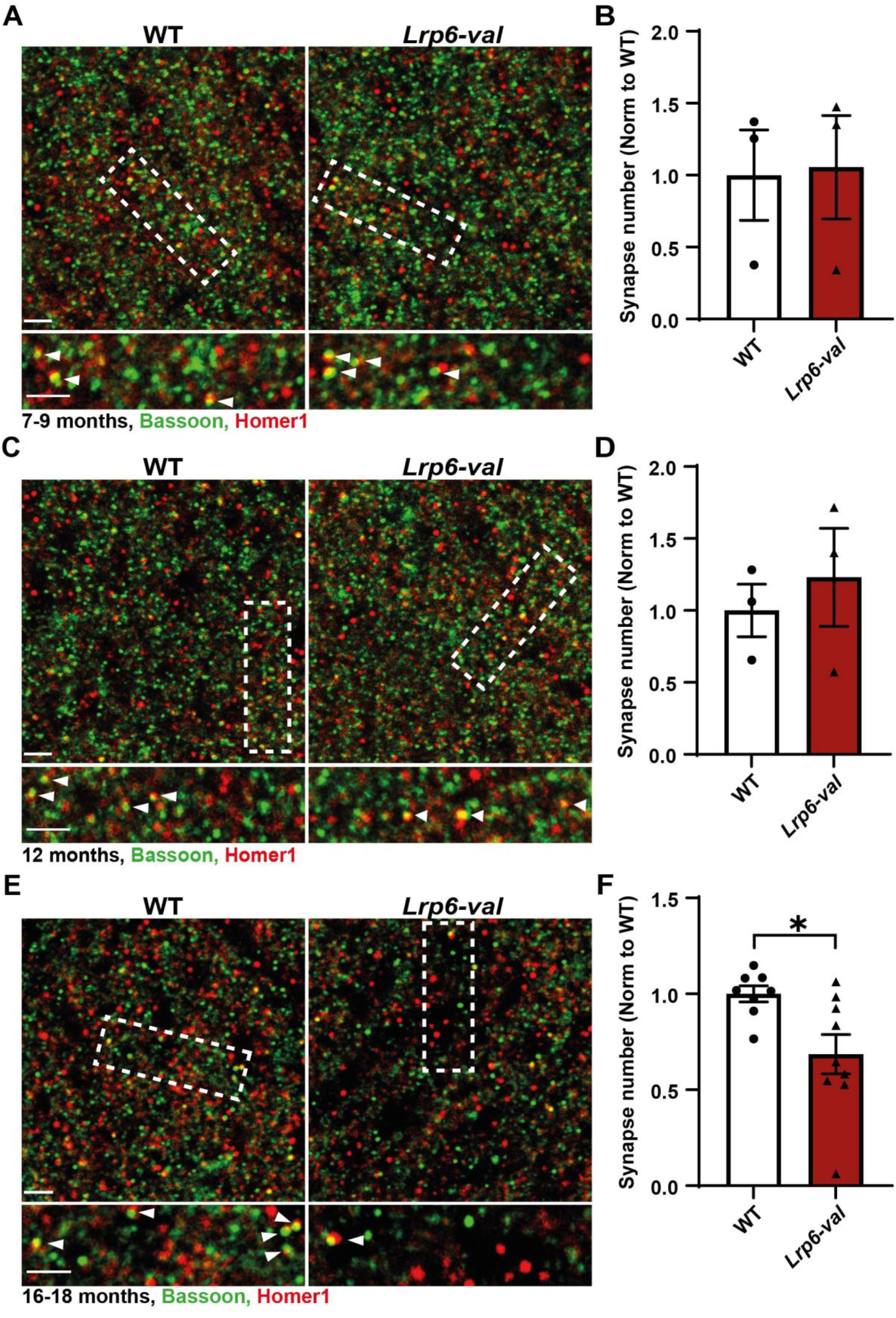
*Lrp6-val* mice exhibit synapse loss with age. A, C, E) Confocal images of the CA1 stratum radiatum of WT and *Lrp6-val* mice labelled with Bassoon (green) and Homer1 (red) at 7-9 months (A), 12 months (C) and 16-18 months (E). Scale bar = 2.5 μm. Insets display high magnification images of synapses. Scale bar = 2 μm. B and D) Quantification of synapse number, based on the co-localisation of pre- and post-synaptic puncta, showed no differences between WT and *Lrp6-val* mice at 7-9 months (B) or 12 months (D). N = 3 per genotype. Unpaired T-test. F) Synapse number was significantly reduced in *Lrp6-val* mice 16-18 months. WT N =8, *Lrp6-val* N = 9. Unpaired T-test. * p < 0.05. Data are represented as mean ± SEM.

### Wnt signalling in impaired in *Lrp6-val* mice and *Lrp6-val* neurons do not respond to Wnt7a

To begin address the molecular mechanisms by which *Lrp6-val* mice exhibit synaptic defects, we investigated if canonical Wnt signalling was affected in aged mice. As a read out, we evaluated the mRNA levels of *Axin2*, a target of canonical Wnt signalling (Jho et al. 2002; Leung et al. 2002) (**Figure S4A**). *Axin2* expression was not affected in 4–7-month-old mice (**Figure S4B**), but it was significantly decreased in 12-15-month-old *Lrp6-val* mice compared to WT (**Figure S4C**). Thus, Wnt signalling is compromised with age *in vivo*, consistent with when we detect synapse defects.

The above findings led us to hypothesize that the LRP6-Val receptor does not signal properly. We therefore interrogated if neurons from *Lrp6-val* mice responded to exogenous Wnts (**Figures 6A**). Previous studies demonstrate that Wnt7a promotes presynaptic assembly (Cerpa et al. 2008; Davis et al. 2008; Sahores et al. 2010) and the formation of excitatory synapses (Ciani et al. 2011) in hippocampal neurons. Consistently, Wnt7a significantly increased the number of synapses in neurons from WT mice compared to neurons exposed to a control vehicle (veh) (**Figures 6B and 6C**). However, Wnt7a was unable to increase synapse number in hippocampal neurons from *Lrp6-val* mice (**Figures 6B and 6C**), indicating that *Lrp6-val* neurons do not respond to Wnt7a. This was not due to decreased levels of the LRP6-Val receptor at the plasma membrane as the surface levels of LRP6 were the same between WT and *Lrp6-val* neurons (**Figures S5A-S5C**). Thus, *Lrp6-val* impairs the ability of neurons to respond to exogenous Wnt7a without affecting its surface localisation.

**Figure 6.**
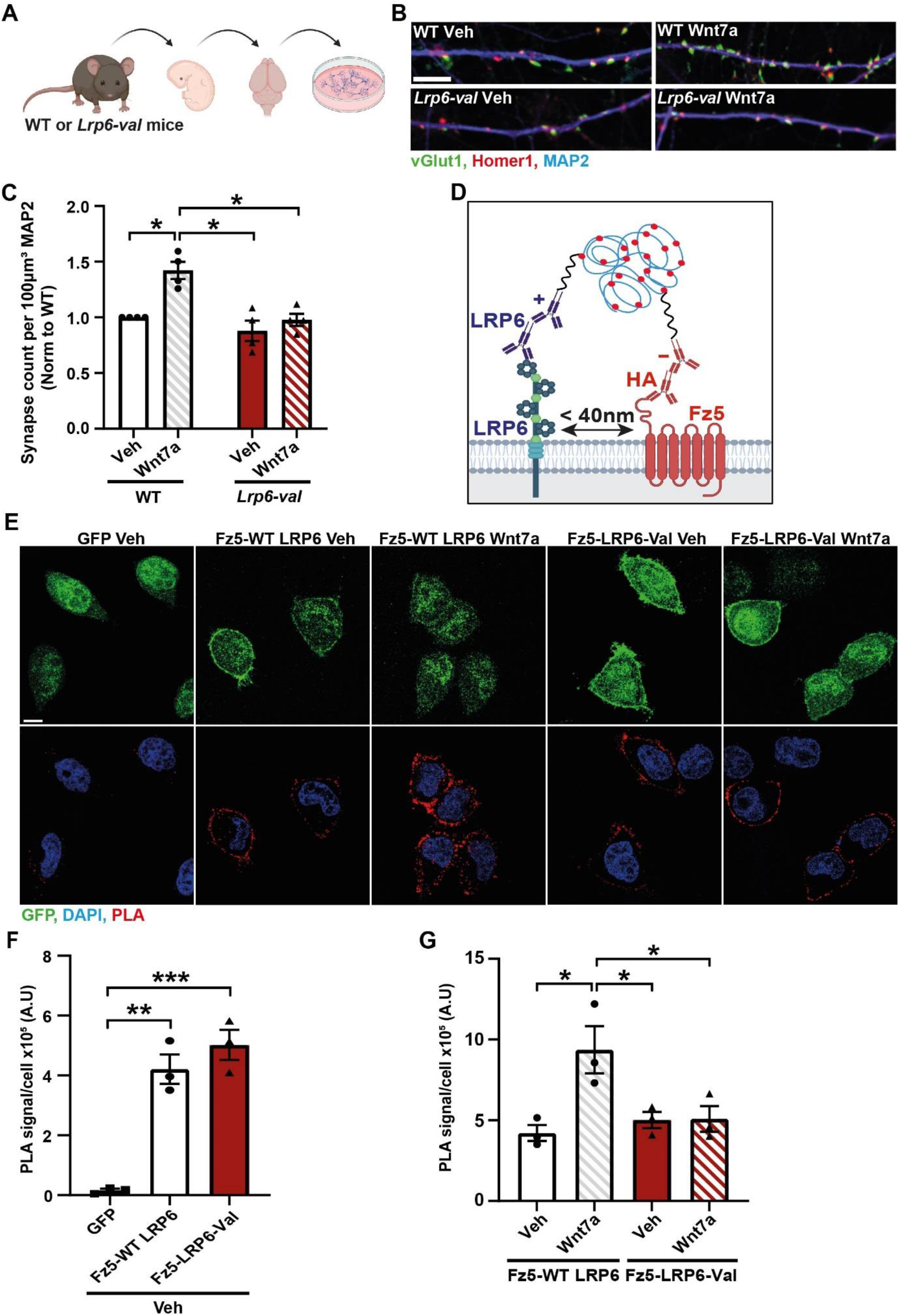
Wnt7a does not stimulate synapse formation in neurons from *Lrp6-val* mice and failed to induce the Wnt receptor complex in cells expressing LRP6-Val. A) Diagram depicting hippocampal neuron isolation from WT and *Lrp6-val* mice. B) Confocal images of WT and *Lrp6-val* hippocampal neurons treated with Wnt7a showing vGlut1 (green), Homer1 (red) and MAP2 (Blue). Scale bar = 5 μm. C) Quantification shows that Wnt7a increased the number of synapses in WT neurons, but Wnt7a had no effect on *Lrp6-val* neurons. N = 4 independent cultures. Two-way-ANOVA with Games-Howell post hoc test. * p < 0.05. D) Schematic of the PLA reaction: antibodies raised from two different species binds to the proteins of interest: LRP6 and Fz5-HA. Secondary antibodies, called PLA probes (+ and −), are linked to specific DNA strands. When proteins are in close proximity (<40 nm), DNA probes can hybridise. PCR amplification with fluorescent complementary probes allows the visualisation of the interaction by fluorescence microscopy. E) Confocal images of HeLa cells expressing GFP (control), WT LRP6 and Fz5-HA or LRP6 Val and Fz5-HA treated with control vehicle (BSA) or Wnt7a. GFP (green), PLA (red) and DAPI (Blue). Scale bar = 10 μm. F) Under basal conditions, the PLA signal intensity per cell was increased in cells expressing WT LRP6 and Fz5-HA or LRP6 Val and Fz5-HA compared to cells only expressing GFP. N = 3 independent experiments. One-way-ANOVA with Tukey’s post-hoc test. ** p < 0.01 and *** p < 0.001. G) Quantification shows that Wnt7a increased the PLA signal in cells expressing WT LRP6 and Fz5-HA but not in cells expressing LRP6 Val and Fz5-HA. N = 3 independent experiments. Two-way-ANOVA with Tukey’s post hoc test. * p < 0.05. Data are represented as mean ± SEM.

### LRP6-Val impairs the Wnt7a-induced LRP6-Fz5 complex formation

The lack of response to Wnt7a in neurons from *Lrp6-val* mice suggested that the presence of LRP6-Val could impair the formation of a complex between LRP6 and Frizzled (Fz) receptors, which is crucial for the activation of the canonical Wnt pathway (Angers & Moon 2009; MacDonald et al. 2009; Hua et al. 2018). We specifically chose to examine the interaction between Fz5 and LRP6 because this Fz receptor is required for Wnt7a-mediated presynaptic assembly in hippocampal neurons (Sahores et al. 2010). We co-expressed Fz5-HA with WT LRP6 or LRP6-Val in HeLa cells to evaluate their interaction, induced by Wnt7a, using proximity ligation assay (PLA), a technique that allows the detection of protein-protein interactions at less than 40nm in intact cells (Söderberg et al. 2006; Klaesson et al. 2018) (**Figure 6D**). The PLA signal was elevated in cells expressing Fz5-HA and WT LRP6 or LRP6-Val when compared to control cells (**Figures 6E and 6F**). No differences in the interaction were observed between cells expressing Fz5-HA and WT LRP6 or Fz5-HA and LRP6-Val, under basal conditions (**Figures 6E, 6F and 6G**). In contrast, Wnt7a significantly increased the intensity of the PLA signal in cells expressing Fz5-HA and WT LRP6 but not in cells expressing Fz5-HA and LRP6-Val (**Figures 6E and 6G**). These results were not due to changes in the surface levels of these receptors, as determined by surface biotinylation (**Figure S5D-S5H**). Together these results demonstrate that the presence of LRP6-Val impairs that formation of the Wnt-induced Wnt receptor complex, which is required for signalling.

### *Lrp6-val* does not affect plaque load in NL-G-F mice

The findings that the *Lrp6-val* variant is associated with late-onset AD (De Ferrari et al. 2007) and that conditional knockout of *Lrp6* in neurons of the APP/PS1 AD mouse model exacerbates the formation of Aβ plaques (Liu et al. 2014), led us to interrogate the impact of the *Lrp6-val* variant on amyloid pathology. We crossed the *Lrp6-val* mice to the *NL-G-F*, a knock-in AD mouse model that carries a humanized Aβ region of APP with three mutations associated with AD (Saito et al. 2014). In *NL-G-F* mice, plaque deposition begins around 2 months, with a significant increase in the number at 7 months (Saito et al. 2014). We assessed the impact of *Lrp6-val* on plaque load in homozygous *NL-G-F* mice. No differences in plaque burden were detected between *NL-G-F* and *NL-G-F;Lrp6-val* mice at 2 months or 7 months (**Figures S6A-S6D**). Furthermore, Aβ coverage (Integrated density) was unaltered in *NL-G-F;Lrp6-val* mice compared to *NL-G-F* mice at 7 months (**Figures S6C and S6D**). Thus, the presence of the *Lrp6-val* variant does not exacerbate plaque load in the *NL-G-F* mice at the ages examined.

Next, we examined the levels of soluble and insoluble Aβ42, as this peptide is the most abundant Aβ species in *NL-G-F* mice (Saito et al. 2014). However, no differences in the level of Aβ42 was observed between *NL-G-F* and *NL-G-F;Lrp6-val* mice at 10 months (**Figure S6E**). These results are consistent with our findings that Aβ plaque number and Aβ coverage are not affected in *NL-G-F* mice.

### *Lrp6-val* exacerbates synapse loss in NL-G-F mice

Although the impact of *Lrp6* cKO on synapse stability has not been examined in the context of AD (Liu et al. 2014), our findings that *Lrp6-val* mice exhibit synaptic defects led us to interrogate the contribution of this SNP to synapse vulnerability in AD. We first evaluated the impact of *Lrp6-val* on synapses in homozygous *NL-G-F* mice at 2 months of age, when plaques begin to form (Saito et al. 2014). However, no differences in synapse number were observed between WT, *Lrp6-val, NL-G-F* or *NL-G-F;Lrp6-val* mice at this early stage (**Figure S7**).

We next investigated the impact of the *Lrp6-val* variant on synapses at 7 months of age in *NL-G-F* mice, when a significant number of plaques are present (Saito et al. 2014), given that synapse loss is particularly pronounced around Aβ plaques (Dong et al. 2007; Koffie et al. 2009; Koffie et al. 2012; Jackson et al. 2016) (**Figure 7A**). Synapse number was quantified at increasing distances from the centre of a plaque, or from a similar area in WT and *Lrp6-val* mice, in the CA1 SR (**Figure 7B**). At 0-10 μm from the centre of a plaque, a significant reduction in synapse number was observed in *NL-G-F* mice when compared to WT mice or to *Lrp6-val* mice (**Figures 7B and 7C**). A further decrease in synapse number was observed between *NL-G-F;Lrp6-val* double mutant mice when compared to *NL-G-F, Lrp6-val* or WT mice. This effect was also observed at further distances from the centre of a plaque (**Figures 7B and 7C**). Thus, carrying the *Lrp6-val* variant exacerbates synapse loss around plaques in *NL-G-F* mice.

**Figure 7.**
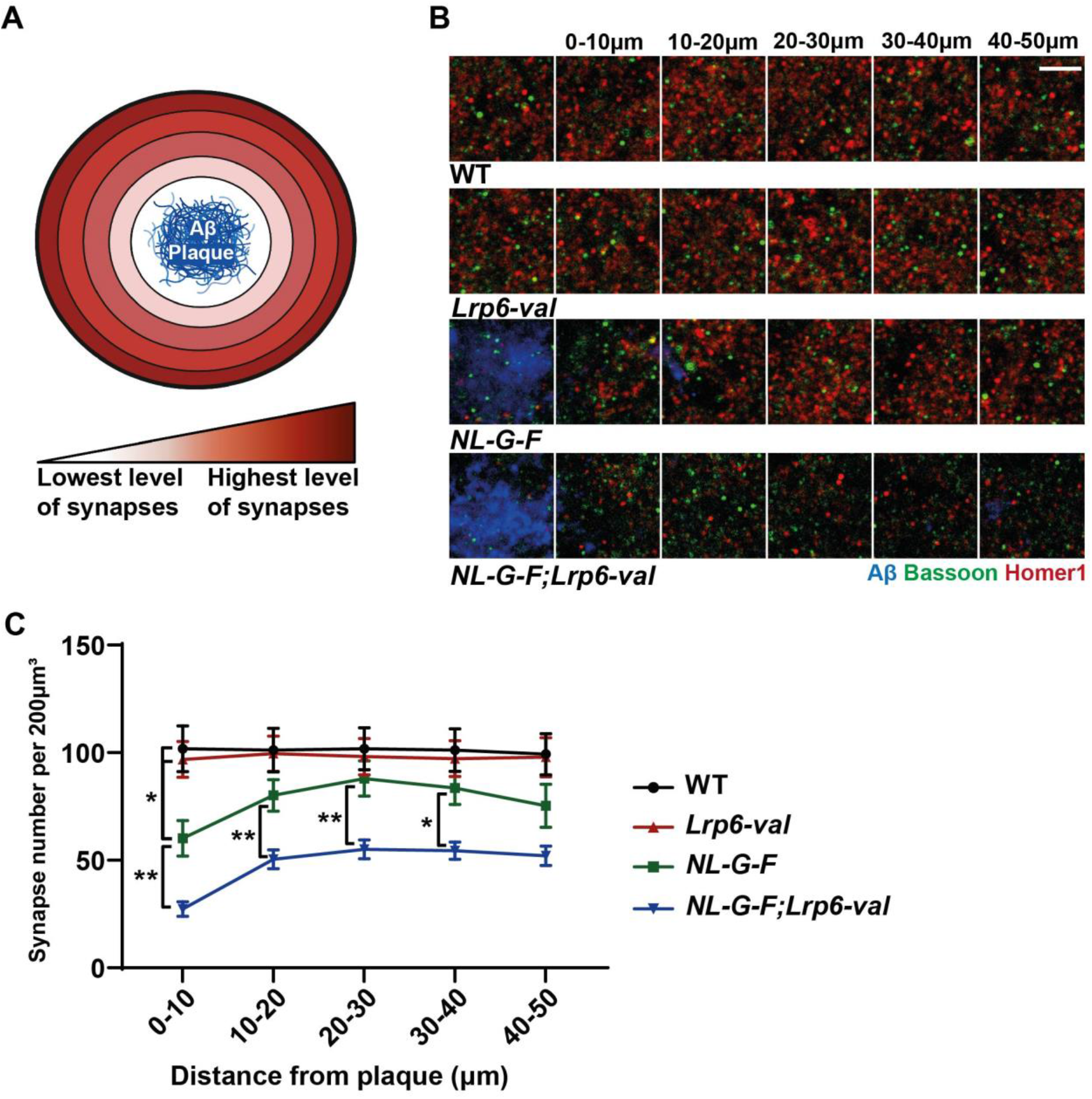
*Lrp6-val* exacerbates synapse loss around plaques in NL-G-F mice at 7 months. A) Diagram shows synapse loss around an Aβ plaque (blue). B) Confocal images of Bassoon (green) and Homer1 (red) at increasing distances from the centre of an Aβ plaque (blue) in *NL-G-F* and *NL-G-F;Lrp6-val* mice, or an equivalent point in WT and *Lrp6-val* mice, in the CA1 stratum radiatum at 7 months. Scale bar = 4 μm. C) *NL-G-F* mice displayed fewer synapses compared to WT mice or to *Lrp6-val* mice at 0-10 μm from the centre of a plaque. Synapse number was reduced in *NL-G-F;Lrp6-val* at 0-40 μm from the center of a plaque compared to *NL-G-F* mice. A significant reduction in synapse number was detected in *NL-G-F;Lrp6-val* when compared to WT mice or *Lrp6-val* mice at all distances from the center of a plaque. N = 6-7 brains per genotype and 2-3 slices per brain. Two-way-ANOVA with Tukey’s post hoc test. * p < 0.05, ** p < 0.01. Data are represented as mean ± SEM.

## Discussion

Here, we evaluate the impact of the *Lrp6-val* variant, linked to LOAD, on synapse connectivity during ageing and in AD. We generated a novel knock-in mouse model carrying this variant. Analyses of homozygous animals reveals that this SNP induces age associated defects in synaptic transmission and neurotransmitter release. *Lrp6-val* mice also exhibit progressive structural synaptic defects in the hippocampus. Importantly, the presence of *Lrp6-val* exacerbates synapse loss around plaques in the *NL-G-F* AD model. Investigation into the molecular mechanisms underlying the synaptic defects elicited by LRP6-Val, reveals a defect in the formation of the LRP6-Fz5 complex mediated by Wnt7a, which explains the decreased downstream signalling that is more evident with age. Our studies also reveal a previously unrecognized molecular mechanism by which this variant impacts Wnt signalling and uncover a role for the *Lrp6-val* variant in synapse degeneration in AD.

The LRP6 receptor, which localises to synapses, promotes the assembly of synapses. Using super-resolution microscopy, we found that LRP6 is present at both the pre- and post-synaptic sites. LRP6 gain-of-function in neurons increases the number of presynaptic puncta and promotes spine growth, consistent with activation of the Wnt pathway in neurons (Davis et al. 2008; Ciani et al. 2011) and in agreement with a previous study showing that LRP6 localizes to both pre- and post-synaptic sites and is required for synaptogenesis in cultured neurons (Sharma et al. 2013). In contrast, expression of the LRP6-Val variant in neurons neither increases the number of presynaptic sites nor affects spine size but induces spine loss. Together, these findings demonstrate that gain-of-function of the LRP6-Val variant in hippocampal neurons fails to promote synapse formation.

Homozygous *Lrp6-val* mice exhibit structural and functional synaptic defects that become more severe with age. At 7-9 months, we observe reduced spine head width without changes in spine number or synapse number, whereas at 12-14 months both spine number and head width and, the number of vGlut1 puncta are decreased. Notably, heterozygous and homozygous *Lrp6-val* mice exhibit a similar spine phenotype (**Figure S8**). Moreover, impairment of synaptic vesicle release, due to a smaller RRP, is exacerbated from 7-9 months to 12-14 months, and basal synaptic transmission is impaired at 12-14 months but not at 7-9 months in homozygous *Lrp6-val*. Although no changes in synapse number are observed at these ages, at 16-18 months, homozygous *Lrp6-val* mice exhibit a reduced number of excitatory synapses. Thus, the presence of the LRP6-Val variant contributes to progressive synapse dysfunction and synapse degeneration.

Multiple variants of *Lrp6* have been linked to various age-associated diseases (Haines et al. 2006; van Meurs et al. 2006; De Ferrari et al. 2007; Sarzani et al. 2011; Wang et al. 2017), including the *Lrp6-val* variant studied in this work, which has an allele frequency of 0.17 in the European population (1000 Genomes Project Consortium et al. 2015). First, *Lrp6-val* is associated with reduced bone mineral density in females over the age of 50 (Riancho et al. 2011) and with a 60% increase in bone fracture risk in older men (van Meurs et al. 2006). Second, *Lrp6-val* is a risk factor for carotid artery atherosclerosis in hypertensive patients over the age of 65 (Sarzani et al. 2011). Finally, *Lrp6-val* is associated with LOAD (De Ferrari et al. 2007). Thus, carrying the LRP6-Val variant results in age-related phenotypes.

What are the molecular mechanisms that contribute to synaptic defects in the *Lrp6-val* mice? The synaptic deficits in *Lrp6-val* mice are not due to defects in the levels or localisation of the LRP6-Val protein, as similar levels are found at synapses and at the cell surface when compared to wildtype LRP6. This suggests possible defects in downstream signalling. Consistent with this hypothesis, Wnt7a fails to induce excitatory synapse formation in neurons isolated from *Lrp6-val* mice. This lack of response correlates with defects in the interaction between Fz5 and LRP6-Val in response to Wnt7a and reduced downstream Wnt signalling as determined by *Axin2* expression at 12-15 months in *Lrp6-val* mice. Moreover, gain-of-function of LRP6-Val attenuates canonical Wnt signalling as evaluated by the TOPFlash assay (De Ferrari et al. 2007). Thus, the presence of LRP6-Val impairs the formation of the Wnt receptor complex in response to Wnt7a, which decreases downstream signalling resulting in synaptic dysfunction and synapse degeneration.

The synaptic defects are manifested with age in the *Lrp6-val* mice. A possible explanation for this phenotype is that canonical Wnt signalling is dampened with age. Indeed, canonical Wnt signalling and several Wnt ligands, including Wnt7a, are reduced in the ageing brain (Hofmann et al. 2014; Bayod et al. 2015; Orellana et al. 2015; Folke et al. 2019). Based on these findings we propose that reduced levels of Wnt proteins with age and the presence of a less functional receptor, such as LRP6-Val, contributes to the manifestation of age-dependent synapse loss in *Lrp6-val* mice.

In the context of AD, our studies demonstrate that carrying the *Lrp6-val* variant increases synapse vulnerability. Indeed, *Lrp6-val* exacerbates synapse loss surrounding plaques in the *NL-G-F* model at 7 months. This is not due to increased Aβ42 levels or plaque load. Our findings are in contrast with those reported using the conditional KO of *Lrp6* crossed to the APP/PS1 AD model where increased Aβ40 and Aβ42 levels and enhancement of plaque load were observed (Liu et al. 2014). These different results could be due to various reasons. First, the cKO of *Lrp6* was studied in APP/PS1, a transgenic model that overexpresses mutant APP and Presenilin1. In contrast, the *NL-G-F* is a knock-in (KI) model with normal levels of APP. Second, our KI model contains a single amino acid substitution in the *Lrp6* gene, which is likely to exhibit a milder phenotype than a cKO model. In summary, our studies demonstrate that a genetic variant of a Wnt receptor, linked to LOAD, increases the risk of synapse loss in AD.

Understanding how risk factors contribute to the pathogenesis of AD is critical for identifying potential therapeutic targets to prevent or ameliorate synapse dysfunction and loss in this condition. Our findings uncover, for the first time, the impact of a genetic variant of *Lrp6* associated with LOAD on synapse vulnerability with age and in the context of AD. Thus, these findings further strengthen the link between deficient Wnt signalling and synapse loss in the ageing and AD brain.

## Acknowledgments

We would like to thank Professors Takashi Saito and Takaomi Saido for the NL-G-F mice, Professor Pico Caroni for the *Thy1-GFP* mice and Professors Xi He, Randall Moon, Bernadette Holdener and Yukiko Goda for DNA constructs. We would like to thank Dr Ian White for performing the EM. We would like to thank to members of our lab for insightful discussions on the data and comments on our manuscript. The MRC (MR/M024083/1, MR/S012125/1 and MC_ST_LMCB_2019) and a Wellcome Trust 4-year Ph.D. scholarship (102267/Z/13/Z) supported our work. Diagrams were created with BioRender.com.

## Authors contribution

P.S. conceived the project. E.M. and K.B. generated the LRP6-Val knock-in mice. J.B. and M.J. contributed to the general characterization of the mice, performed cell biology, imaging and biochemical experiments. J.B. and T. D. performed the electrophysiology experiments. T.D. performed biochemical experiments. A.G. advised on the design and interpretation of electrophysiological experiments. All authors participated in the design of experiments, interpretation of data and writing of the manuscript.

## Materials and Methods

### Animals

Experiments with mice were carried out under personal and project licenses granted by the UK Home Office in accordance with the Animals (Scientific Procedures) Act 1986. Animals were housed in ventilated cages with a 12-hour light-dark cycle and *ad libitum* access to food and water. Both male and female animals were used. Ages are specified in the figure legends.

### Generation of LRP6-Val mutant mice

Single-strand oligonucleotides (ssODNs) were synthesised by Integrated DNA Technologies (IDT)(ssODN:CATCAGAGGCAGTCTCAGGCTGTGGCTTTGGAACATACCCTTTCTCGGGGTTTACCACAACGGCTCAGGTCTGTCTTGCTCGCCTTTTAGAACCACTCCAACTGATCGTCCATCTAATC). ssODNs were positioned adjacent to the gRNA site: ACAGACCTCGAGCCATTGTGG. gRNA oligonucleotides were synthesised, the two strands annealed and cloned using BsaI into a vector containing the gRNA backbone and a T7 promoter for RNA production. For Cas9 mRNA production, the vector from (Cong et al. 2013), was modified to contain the T7 promoter. 4-5-week-old C57BL/6NTac females were super-ovulated by injection of 5 IU of pregnant mare’s serum (PMSG) and 48 hours later, 5 IU of human chorionic gonadotrophin (HCG) was injected. Females were mated with C57BL/6NTac males. Cumulus oocyte complexes were dissected from oviducts 21-22 hrs post HCG and treated with hyaluronidase. Fertilised 1-cell embryos were maintained at 37°C in KSOM media prior to cytoplasmic injection. 24-27 hrs post HCG, 50 ng/μL Cas9 mRNA, 25 ng/μL gRNA (each) and 100 ng/μL oligonucleotide were injected into the cytoplasm of fertilised 1-cell embryos held in FHM medium. Viable embryos were transferred on the same day by oviducal embryo transfer into 0.5 days post-coital pseudo-pregnant female F1 (CBA/C57BL/6J) recipients. Homozygous *Lrp6-Val* C57BL/6NTac mice were backcrossed to C57BL/6J mice.

### Genotyping

*Lrp6-val* mice were crossed to the *Thy1-GFP* mice or *NL-G-F* model to obtain *Lrp6-val;Thy1-GFP* mice and *NL-G-F;Lrp6-val* mice respectively. Genotyping was performed on ear biopsies using the following primers: *Lrp6* WT Forward: GATACGTTGCTTTAATGCCTTTAGCAAGACAGACCTCGAGCAA, *Lrp6-val* Forward: TGGCGGCAAGACAGACCTCGAGCAG, *Lrp6* (WT and Val) Reverse: AACGCGCAACGAAGGGTGAGGAGGCATCA. *NL-G-F:* 5’-ATCTCGGAAGTGAAGATG-3’, 5’-ATCTCGGAAGTGAATCTA-3’, 5’-TGTAGATGAGAACTTAAC-3’ and 5’-CGTATAATGTATGCTATACGAAG-3’ (Saito et al. 2014). GFP: forward 5’-TCTGAGTGGCAAAGGACCTTAGG-3’ and reverse 5’-CGCTGAACTTGTGGCCGTTTACG-3’ (Feng et al. 2000).

### Hippocampal culture and transfection

Rat hippocampal cultures were prepared from embryonic day 18 (E18) embryos from Sprague-Dawley rats as previously described (Dotti et al. 1988). Cultures were maintained until 13-14 days in vitro (DIV) to presynaptic terminals or 21 DIV to investigate dendritic spines.

Mouse hippocampal neurons were prepared from E15.5-E16.6 WT or *Lrp6-val* mice and maintained until 12-21 DIV. Neurons were treated at with 200 ng/ml recombinant Wnt7a (PeproTech, 120-31) or BSA (control vehicle) for 3 hours at 12 DIV for analyses of synapses.

Rat hippocampal neurons were transfected using calcium phosphate transfection method at 7-9 DIV with DNA constructs expressing EGFP-actin, human LRP6 WT or human LRP6-Val and MESD (mesoderm development LRP chaperone protein), which is required for maturation and trafficking of LRP6 (Hsieh et al. 2003; Li et al. 2006; Liu et al. 2009). Control neurons were transfected with EGFP-actin.

### HeLa cell culture and transfection

HeLa cells were grown in DMEM (Gibco) supplemented with 10% FBS and 1% penicillin/streptomycin (Gibco) and maintained at 37°C and 5% CO_2_. Cells were seeded on 12mm glass coverslips at a density of 13.4×10^4^ cells/cm^2^ for PLA assays and at 18.6×10^4^ cells/cm^2^ for surface biotinylation.

Cells were transfected with EGFP, Fz5-HA, MESD and LRP6 WT or LRP6-Val plasmids, using Lipofectamine 3000 (Invitrogen) and OPTIMEM according to the manufacturer’s protocol, for 3hrs. The transfection medium was then replaced with OPTIMEM. 48hr post transfection, cells were treated with serum free DMEM containing recombinant Wnt7a (200ng/ml, PreproTech, 120-31) or BSA (control) for 30 minutes.

### List of plasmids

LRP6 WT (Addgene plasmid # 27242) and Fz5-HA were gifts from Professor Xi He. MESD-Flag was a gift from Professor B. Holdener. EGFP-actin was a gift from Dr Y. Goda. LRP6 Val-HA was a gift from Professor R. T. Moon. The untagged LRP6-Val plasmid used in this paper was generated using constructs kindly provided by Professor’s R. T. Moon and Xi He.

### Brain section preparation

For cryostat sections used for immunostaining, 4% paraformaldehyde (PFA) fixed brains were immersed in 30% sucrose and frozen in precooled 2-methylbutane. 30-50μm sagittal sections were cut using a cryostat and stored at −20°C. Vibratome sections were prepared as previously described (McLeod et al. 2017). Briefly, brains were rapidly dissected, immersed in artificial cerebrospinal fluid (aCSF) and 250-300μm thick sagittal brain slices were cut. Then, slices were fixed in 4% PFA/ 4% sucrose in PBS.

### Immunofluorescence staining

Immunofluorescence staining was performed as previously described (McLeod et al. 2017). Slices were permeabilized and blocked in 0.5% Triton-X-100 + 10% donkey serum in PBS for 3-4 hours at room temperature and then incubated with primary antibodies overnight at 4°C.

Secondary antibodies (1:500; Alexa Fluor, Invitrogen) were incubated for 2 hours at room temperature. Slices were incubated in DAPI, washed with PBS and mounted with Fluoromount-G (SouthernBiotech).

Cultured hippocampal neurons were fixed in 4% PFA with 4% sucrose in PBS for 20 minutes at room temperature. Neurons were permeabilized in 0.05% Triton-X-100 in PBS for 5 minutes, blocked in 5% BSA for 1 hour, both at room temperature, and the incubated with primary antibodies overnight at 4°C. Secondary antibodies (1:600; Alexa Fluor, Invitrogen) were incubated for 1 hour at room temperature. Neurons were incubated with DAPI, washed with PBS and mounted with FluorSave (Millipore).

### Proximity ligation assay (PLA)

Proximity ligation assay was performed according to the manufacturer’s protocol (Sigma Aldrich). Briefly, cells were washed with PBS, fixed with warm 4% PFA for 15 minutes then blocked for 60 minutes at 37°C with the Duolink Blocking solution. Anti-LRP6 (R&D, AF1505) and anti-HA (Sigma, H6908) primary antibodies in Duolink Antibody diluent solution (1:800) were incubated overnight at 4°C. Cells were washed three times with the Duolink Wash Buffer A and incubated with the anti-rabbit MINUS and anti-goat PLUS PLA probes in Duolink Antibody diluent solution for 1 hour at 37°C. After two washes in the Wash Buffer A, cells were incubated with the Duolink Ligation solution for 30 minutes at 37°C. After two Wash Buffer A washes, cells were incubated with the Duolink Amplification solution for 1 hour 40 minutes at 37°C. Cells were washed 2×10 minutes in Wash Buffer B and for 1 minute in 0.01x Wash Buffer B. Cells were then permeabilised for 10 minutes with PBS 0.1% Triton and blocked in 5% BSA for 1 hour. Anti-GFP (1:500, Millipore, 06-896) primary antibody was added for 1 hour at RT, followed by 3 PBS washes and the addition of AlexaFluor 488 chicken for 1 hour at RT. After 3x PBS washes, coverslips were mounted using 5μl per coverslip of Duolink In Situ Mounting Medium with DAPI.

### List of primary antibodies

APP (6E10) (Novus Biotech, NBP2-62566), Amyloid-β (BioLegend, 803001), β-Actin (Cell Signalling Technology, 4970), Bassoon (Novus Biologicals, NB120-13249), GFP (Millipore, 06–896), GFP (Invitrogen, A-6455), Homer1, (Synaptic systems, 160002), Homer1, (Synaptic systems, 160003), LRP6 (Abcam, ab134146), LRP6 (R&D, AF1505), LRP6 (Cell Signalling Technology, 2560), LRP6 (Cell Signalling Technology, 3395), HA (Sigma, H6908), HA (Roche, 11867423001), MAP2 (Abcam, ab5392), MAP2 (Abcam, ab92434), NeuN, (Cell Signalling Technology, 12943), PSD-95 (Millipore, MAB1598), α-Tubulin (Sigma, T9026), vGlut1 (Millipore, AB5905), Vinculin (Sigma, V4505).

### Confocal microscopy

Images were acquired on a Leica SP8 or an Olympus FV1000 inverted confocal microscope. For analysis of synaptic puncta and dendritic spines in brain sections, 3 images (stacks) were acquired per brain section and 3 brain sections were analysed per animal. For hippocampal cultures, 8-13 images (stacks) were acquired per condition. Each stack comprised 8-11 equidistant planes, 0.25μm apart, and were acquired using a 63× 1.40 NA oil objective. For analysis of NeuN staining, a stack of 31 equidistant planes, 0.5μm apart, was acquired for each brain section with a 10× 0.40 NA objective on a Leica SP8. For plaque analysis of 2 month-old-mice, images were acquired on a Leica SP5. For each brain section, one stack of 25 equidistant planes, 0.99μm apart, was acquired using a 10x 0.30 NA objective. For analysis of plaques in 7 month-old-mice, a tile scan per brain section was acquired with a 20x 0.75 NA objective on a Leica SP8. Each stack comprised 59 equidistant planes 0.35μm apart. For the PLA experiments in HeLa cells, stacks of 10 equidistant planes with a z step of 0.5μm were acquired using a 40x oil objective on a Leica SP8. Two coverslips per experimental condition were imaged with 4-5 images per coverslip acquired.

### SIM

Structured Illumination Microscopy (SIM) was performed on a Zeiss Elyra S.1 microscope with a 63x oil-immersion objective (NA 1.40). A z-stack of 15-20 equidistant planes were acquired. For each field of view, nine images were acquired, using three different rotations and phases of structured illumination (a grid pattern) on the sample.

### EM

Sagittal brain sections of 200 μm thickness, were cut on a vibratome then fixed in 2 % PFA/2.5 % Glutaraldehyde solution, overnight at 4°C. Samples were post-fixed in 1 % O_s_O_4_ for 1 hr at 4°C and stained with 1 % thiocarbohydrazide for 20 minutes at RT, 2 % OsO4 for 30mins at RT, 1 % uranyl acetate (UA) ON at 4°C and lead aspartate for 30 minutes at 60°C. Next, slices were dehydrated in graded alcohol and embedded in resin. Ultra-thin sections (70 nm) were then cut using a diamond ultra 45-degree knife (Diatome) on a Leica UC7 ultramicrotome and collected on 2 x 1 mm slot grids. Images were acquired on a transmission electron microscope (T12 Tecnai Spirit Bio-Twin, FEI) each covering 5.8 μm^2^ at 0.77 nm/pixel.

### Image analysis

Image analyses were performed using Volocity software (Perkin Elmer). For hippocampal cultures and brain sections, customized thresholding protocols were used to detect pre- and postsynaptic puncta. Synapses were quantified as co-localized pre-and postsynaptic puncta. Analysis of dendrite spines was performed blind to the genotype or treatment. Dendritic spines were measured manually along 3-4 sections of dendrite. Spine size was quantified using the line tool and measuring the maximum spine head width. For the PLA experiment, only transfected cells were selected (based on GFP signal) and the total PLA signal intensity was quantified and divided by the number of cells analysed. For SIM images, synapses were identified manually and the co-localisation of LRP6 with synaptic markers was determined using Volocity. For neuronal number, using Volocity the number of NeuN and DAPI positive cells were quantified and divided by the total number of DAPI positive cells. In *NL-G-F* and *NL-G-F;Lrp6-val* mice, plaques were manually counted at 2 months, blind to the genotype. At 7 months, plaque analyses were performed, blind to the genotype, using Fiji as previously described (Pickett et al. 2019). Images were thresholded and then the % area was measured. The particle analysis tool was used to count the number of plaques. For EM images, synaptic vesicles were manually counted within 200 nm of the active zone in high-magnification images. PSD length was quantified using a line tool in ImageJ. Vesicle number was normalised to PSD length.

### Electrophysiological Recordings

Electrophysiological recordings were performed in 7-8 month-old and in 12-13-month-old male and female mice. Acute transverse hippocampal slices (300 μm thick) of WT control mice and homozygous *Lrp6-val* mice were cut with a Leica VT-1000 vibratome in ice-cold artificial cerebrospinal fluid (ACSF) bubbled with 95% O_2_/5% CO_2_ containing: NaCl (125 mM), KCl (2.4 mM), NaHCO_3_ (26 mM), NaH_2_PO_4_ (1.4 mM), D-(+)-Glucose (20 mM), CaCl_2_ (0.5 mM) and MgCl_2_ (3 mM). Brain slices from 12-13 months old mice were then transferred for 5 minutes into a series of three different oxygenated (95% O_2_/5% CO_2_) chambers in the same ACSF base but with a gradual temperature and component variations: 1) 21 °C, MgCl_2_ (1 mM) and CaCl_2_ (0.5 mM) then placed at 36 °C for 5 minutes; 2) 36 °C, MgCl_2_ (1 mM) and CaCl_2_ (1 mM); and 3) 36 °C with MgCl_2_ (1 mM) and CaCl_2_ (2 mM) before cooling to 21 °C for at least 1 hour before recordings. Brain slices were placed in a chamber on an upright microscope and constantly perfused with NaCl (125 mM), KCl (2.4 mM), NaHCO_3_ (26 mM), NaH_2_PO_4_ (1.4 mM), D-(+)-Glucose (20 mM), MgCl_2_ (1 mM) and CaCl_2_ (2 mM) supplemented with 10 μM bicuculline, to block GABA currents, at room temperature.

Whole-cell patch-clamp recordings were made from pyramidal cells in the CA1 region voltage-clamped at −60mV using patch pipettes with a resistance of 4-8 MΩ when filled with a caesium gluconate pipette solution composed of: D-gluconic acid lactone (130 mM), Hepes (10 mM), EGTA (10 mM), NaCl (10 mM), CaCl_2_ (0.5 mM), MgCl_2_ (1 mM), ATP (1 mM) and GTP (0.5 mM), QX314 (5 mM) adjusted to pH 7.2 with CsOH. To evoke postsynaptic currents, a bipolar concentric stimulating electrode (FHC) connected to a Grass S48 stimulator was placed around 100-200 μm from the patched cell. Cell input-output recordings were made with the stimulus pulse varied between 9 and 50V with a pulse width of 0.1 ms and delivered at a rate of 0.1 Hz. At least three responses per stimulation intensity were averaged per cell. Paired pulse ratio (PPR) stimuli were delivered at a rate of 0.2 Hz with varying inter-stimulus intervals, ranging from 50 ms to 200 ms. Stimulus intensity was adjusted for each cell to elicit ~50% of the maximal response. PPR was calculated as the ratio of the peak amplitude of the second response over the first response and at least seven responses were averaged per cell for each inter-stimulus interval. For RRP size, initial fusion efficiency, and SV recycling rate calculation, CA1 cell EPSCs were recorded in response to 3 s duration trains of stimulation at 20 Hz and estimated as previously described (Wesseling & Lo 2002; Ciani et al. 2015).

The following two equations were used to estimate RRP size, fusion efficiency *fe*), and vesicle recycling rate (α) using cumulative charge (Wesseling & Lo 2002; Stevens & Williams 2007) in analysis of 20 Hz stimulus-evoked trains of EPSCs:

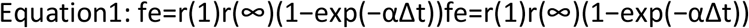

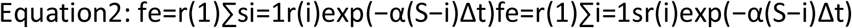

*r*(1) is the charge of the first EPSC in the train, *r(i*) is the charge passed by the *i*th EPSC, *r*(∞) was calculated from the average charge of the last 10 EPSCs in the train, and Δ*t* is the stimulus interval in the train. The RRP was estimated as RRP =*r*(1)/*fe*.

Currents were recorded using an Axopatch 200B amplifier and low-pass filtered at 1 kHz and digitized at 10 kHz. Online monitoring of the data was performed using WinEDR and offline analysis using both WinEDR and WinWCP software (freely available online at http://spider.science.strath.ac.uk/sipbs/software_ses.htm).

### Surface biotinylation and Western Blots

Surface biotinylation was performed on using Sulfo-NHS-LC-LC biotin (Thermo, EZ-Link Sulfo NHS-LC-LC-Biotin) and streptavidin-agarose beads (Thermo Fisher). Samples were run on an 8% SDS-PAGE gel. Hippocampal homogenates from WT and *Lrp6-val* mice at 4 months old were resolved on 8-12 % SDS-PAGE gels.

### ELISA

*NL-G-F* and *NL-G-F;Lrp6-val* hippocampal tissue was homogenized in RIPA, followed by guanidine hydrochloride (GuHCL). Aβ42 peptides were quantified using the human Aβ 1-42 ELISA kit (WAKO), following the manufacturer’s instructions.

### Quantitative PCR (qPCR)

RNA was extracted from frozen hippocampal tissue using Trizol (Life Technologies) and the Direct-zol RNA MiniPrep Kit (Zymo Research), following the manufacturer’s instructions. First-strand cDNA synthesis was performed with the RevertAid H Minus First Strand cDNA Synthesis kit (Thermo Fisher Scientific), following manufacturer’s instructions. Quantitative PCR (qPCR) was performed using GoTaq qPCR Master Mix (Promega). The primers used for *Lrp6* were: forward 5’-TCTTGTGGTTGTCTGGTGTGGAG-3’ and reverse 5’-AGAAGACATATCAGAAAATGCAGGAGG-3’. The primers used for *Axin2* were: forward ‘5-GAGGGACAGGAACCACTCG-3’ and reverse: ‘5-TGCCAGTTTCTTTGGCTCTT-3’.

### Statistical analyses

All results are presented as mean ± SEM. Statistical analyses were performed in GraphPad Prism (Version 9) or SPSS (IBM, Version 27). Normality was assessed with Shapiro-Wilk or Kolmogorov-Smirnov tests. Normally distributed data was analyzed using T-tests for two conditions, one-way ANOVA for two or more conditions or two-way ANOVA for experiments with two independent variables. Post hoc tests are detailed in the figure legends. Non-normally distributed data was analyzed with non-parametric tests such as Kruskal-Wallis or Mann-Whitney. Statistical significance was accepted as * p < 0.05, ** p < 0.01, *** p < 0.001.

