## Supplementary Materials for "A genetic variant of the Wnt receptor LRP6 accelerates synapse degeneration during ageing and in Alzheimer’s disease"

* Corresponding author

#### Supplementary Figures

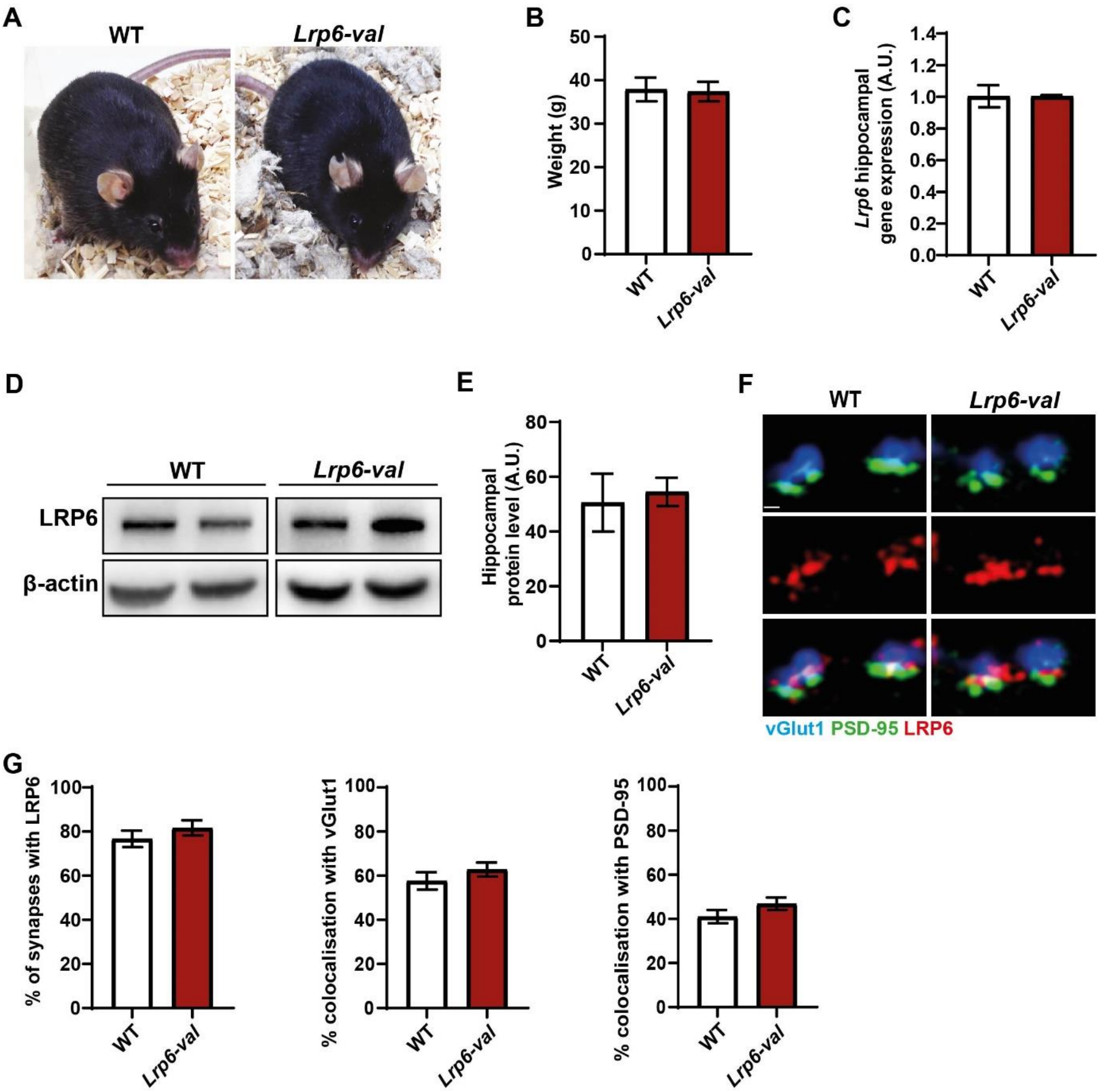


**Figure S1**

**Figure S1. Characterisation of homozygous *Lrp6-val* knock-in mice**. A) Images of WT and *Lrp6-val* mice at 7 months showed that these mice developed normally and have no visible external abnormalities. B) Adult *Lrp6-val* mice had the same weights as control WT mice. Weights of male mice were measured at 4-8 months of age. WT N = 9. *Lrp6-val* N = 10. Unpaired T-test. C) Quantitative RT-PCR analyses of *Lrp6* mRNA levels in the hippocampus of WT and *Lrp6-val* mice at 3-4 months of age showed no changes the expression of *Lrp6*. WT N = 5, *Lrp6-val* N = 6. Unpaired T-Test. D) Hippocampal LRP6 protein levels in WT and *Lrp6-val* mice at 4 months. E) No changes in LRP6 levels (normalised to β-actin) were observed between WT or *Lrp6-val* mice. N = 3 per genotype. Unpaired T-test. F) SIM images of excitatory synapses containing LRP6 (red) from WT and *Lrp6-val* hippocampal neurons showing vGlut1 (blue) and PSD-95 (green). Scale bar = 0.2 µm. G) LRP6 receptor exhibited similar pre- and post-synaptic localisation in WT and *Lrp6-val* mice. 2 independent cultures, 6-8 images per culture. Unpaired T-tests. Data are represented as mean ± SEM.

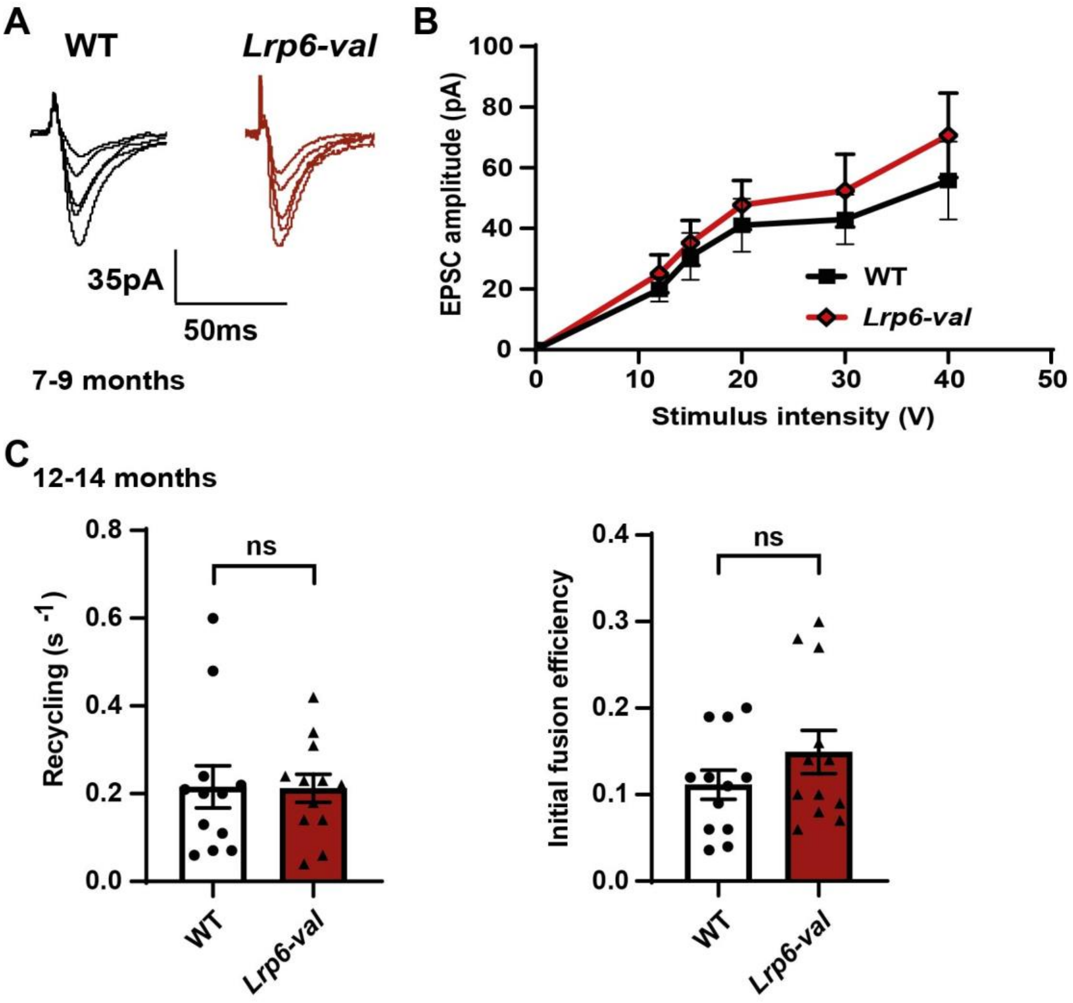


Figure S2

**Figure S2. Basal synaptic transmission is unaffected in *Lrp6-val* mice at 7-9 months and vesicle recycling and initial fusion efficiency are unchanged in *Lrp6-val* mice at 12-14 months.** A) Representative traces of post-synaptic currents elicited at different stimulation intensities. B) No differences were detected in the input-output curves at 7-9 months. N = 12-14 cells recorded from 4-5 animals per genotype. Repeated-measures one-way-ANOVA. C) Graphs display the recycling rate and initial fusion efficiency obtained from all cells. No differences were observed in *Lrp6-val* mice when compared to WT mice at 12-14 months. *N* = 12 cells from 4 animals per genotype. Unpaired Student's *T-t*est. Data are represented as mean $\pm$ SEM.

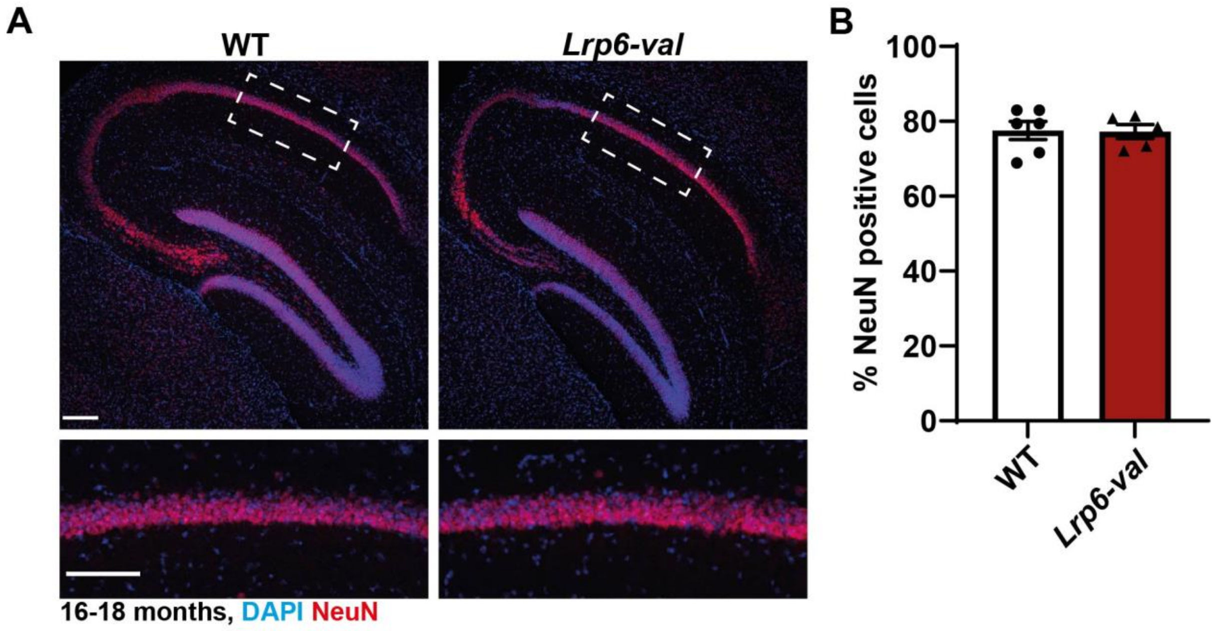


**Figure S3**

**Figure S3. *Lrp6-val* mice do not display neuronal loss at 16-18 months.** A) Confocal images of the hippocampus of WT and *Lrp6-val* mice labelled with DAPI (blue) and NeuN (Red). Scale bar = 150 µm. Insets show higher magnification images of the CA1 region. Scale bar = 100 µm. B) Quantification revealed no differences in the percentage of NeuN positive cells between WT and *Lrp6-val* mice. WT N = 6, *Lrp6-val* N = 5. Unpaired T-test. Data are represented as mean $\pm$ SEM.

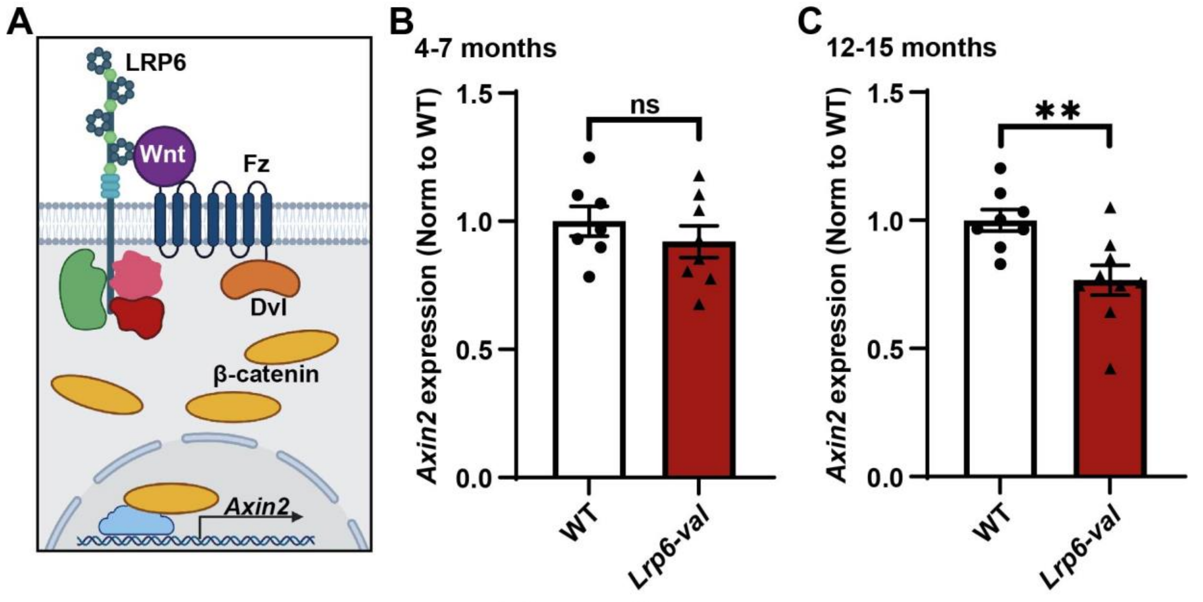


**Figure S4**

**Figure S4. *Axin2* mRNA levels are reduced at 12-15 months in *Lrp6-val* mice.** A) Schematic of the Wnt signalling pathway showing regulation of *Axin2* expression. B and C) Quantitative RT-PCR analyses of *Axin2* expression in the hippocampus of WT and *Lrp6-val* mice at 4-7 months (B) and 12-15 months (C). *Axin2* expression was reduced in *Lrp6-val* mice at 12-15 months but not before. 4-7 months: WT N = 7, *Lrp6-val* N = 8. 12-15 months: WT N = 8, *Lrp6-val* N = 9. Unpaired T-tests. ** $p < 0.01$. Data are represented as mean $\pm$ SEM.

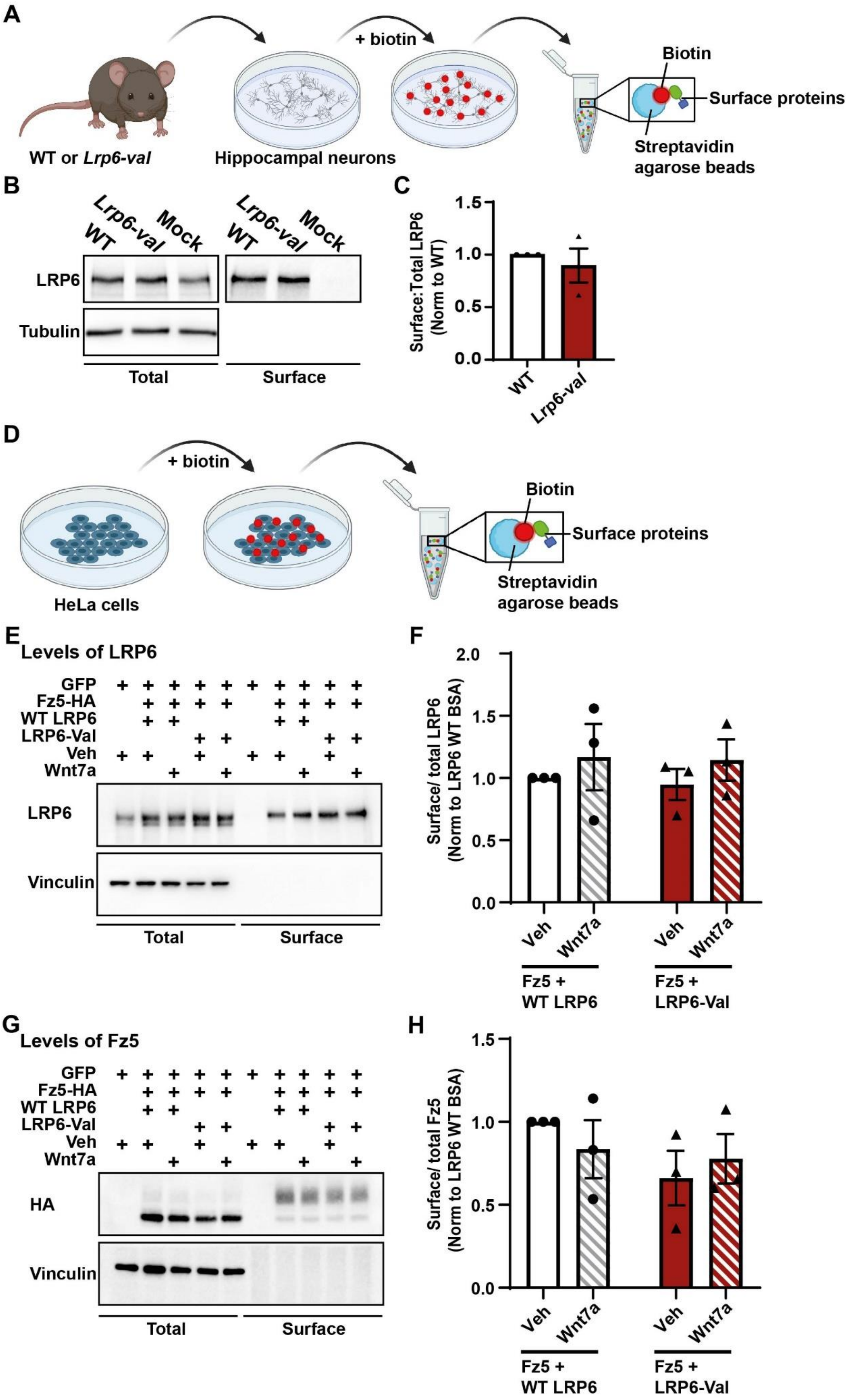
A
+ biotin
Biotin
Surface proteins
WT or Lrp6-val
Hippocampal neurons
Streptavidin agarose beads
B
WT
Lrp6-val
Mock
LRP6
Tubulin
Total
Surface
C
Surface:Total LRP6 (Norm to WT)
1.5
1.0
0.5
0.0
WT
Lrp6-val
D
+ biotin
Biotin
Surface proteins
HeLa cells
Streptavidin agarose beads
E
Levels of LRP6
GFP
Fz5-HA
WT LRP6
LRP6-Val
Veh
Wnt7a
LRP6
Vinculin
Total
Surface
F
Surface/ total LRP6 (Norm to LRP6 WT BSA)
2.0
1.5
1.0
0.5
0.0
Veh
Wnt7a
Fz5 + WT LRP6
Fz5 + LRP6-Val
G
Levels of Fz5
GFP
Fz5-HA
WT LRP6
LRP6-Val
Veh
Wnt7a
HA
Vinculin
Total
Surface
H
Surface/ total Fz5 (Norm to LRP6 WT BSA)
1.5
1.0
0.5
0.0
Veh
Wnt7a
Fz5 + WT LRP6
Fz5 + LRP6-Val

**Figure S5**

**Figure S5. Surface localisation of LRP6 is not affected in *Lrp6-val* neurons or or cells expressing LRP6-Val.** A) Diagram of surface biotinylation experiments performed in hippocampal neurons from WT and homozygous *Lrp6-val* mice. Neurons were incubated with biotin (red circles). Biotin bound surface proteins (Green and blue shapes) were pulled down with streptavidin-agarose beads (blue circles). B) Surface biotinylation analyses of LRP6 were performed on neurons isolated from WT and homozygous *Lrp6-val* mice. C) No differences in the ratio of surface to total LRP6 were observed. N = 3 independent cultures. Mann-Whitney. D) Schematic of surface biotinylation analyses of HeLa cells. E) Western blot analyses of LRP6 levels following surface biotinylation of HeLa cells expressing Fz5-HA and WT LRP6 or LRP6-Val and treated with Wnt7a. F) No differences in the ratio of surface to total LRP6 were observed after Wnt7a treatment. N = 3 independent cultures. Two-way-ANOVA with Games-Howell post hoc test. G) Western blot analysis of Fz5-HA following surface biotinylation of HeLa cells expressing Fz5-HA and WT LRP6 or LRP6-Val and treated with Wnt7a. We observed a higher molecular weight of Fz5-HA at the surface, which is probably due to changes in glycosylation of this receptor (Koo et al. 2012). H) The ratio of surface to total Fz5-HA were unchanged after Wnt7a treatment. Both bands observed for HA were quantified. N = 3 independent cultures. Two-way-ANOVA with Games-Howell post hoc test. Data are represented as mean $\pm$ SEM.

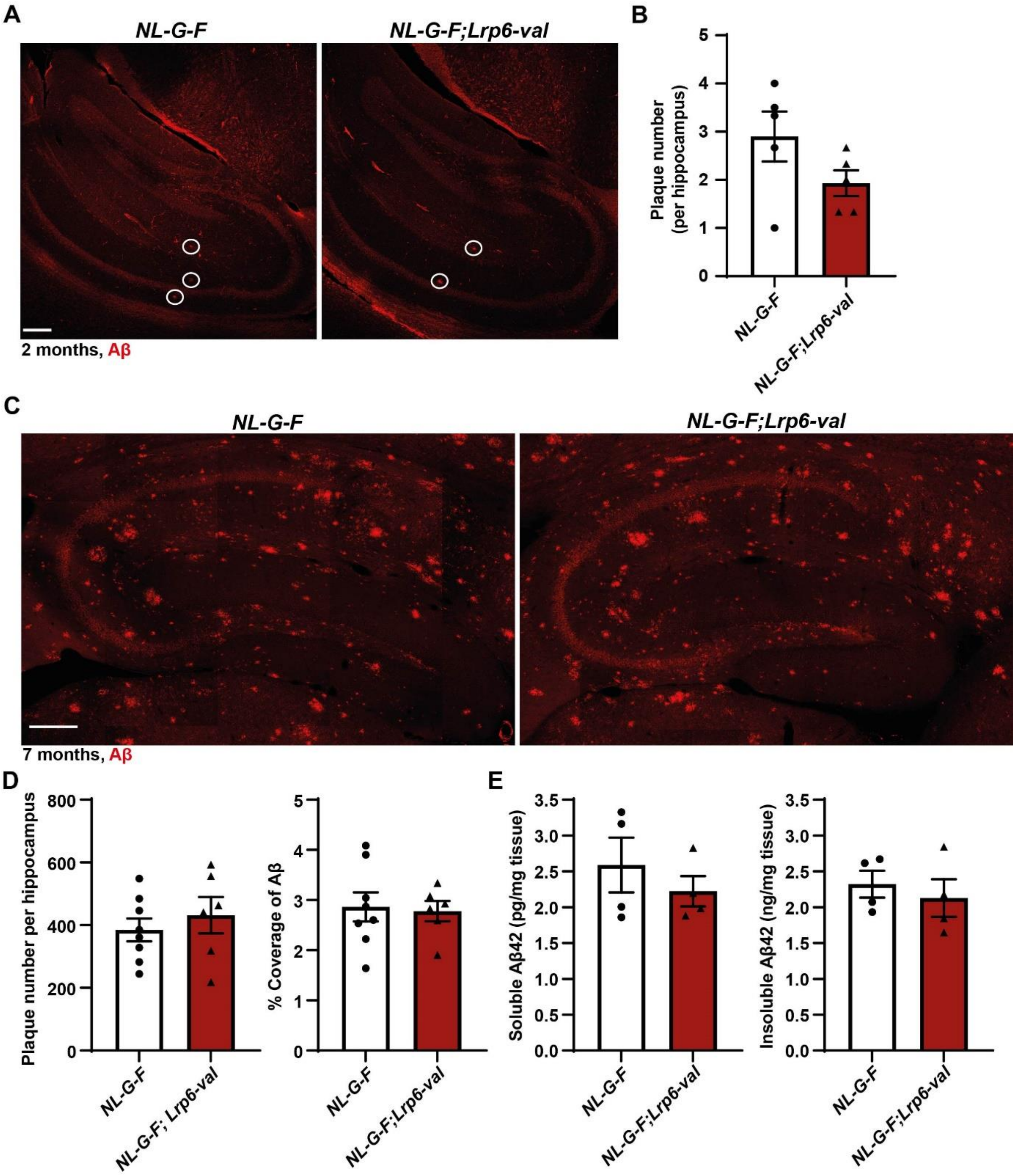
A
NL-G-F
NL-G-F;Lrp6-val
2 months, Aβ
B
Plaque number (per hippocampus)
5
4
3
2
1
0
NL-G-F
NL-G-F;Lrp6-val
C
NL-G-F
NL-G-F;Lrp6-val
7 months, Aβ
D
Plaque number per hippocampus
800
600
400
200
0
NL-G-F
NL-G-F; Lrp6-val
% Coverage of Aβ
NL-G-F;Lrp6-val
E
Soluble Aβ42 (pg/mg tissue)
3.5
3.0
2.5
2.0
1.5
1.0
0.5
0.0
Insoluble Aβ42 (ng/mg tissue)

**Figure S6**

**Figure S6. Plaque load and Aβ42 levels are unaltered in *NL-G-F;Lrp6-val* mice.** A) Confocal images of Aβ (red) in the hippocampi of *NL-G-F* and *NL-G-F;Lrp6-val* mice at 2 months. Scale bar = 150 µm. B) Quantification shows no differences in plaque number were detected. *NL-G-F* N = 5, *NL-G-F;Lrp6-val* N = 5. Unpaired T-test. C) Aβ plaques (red) in the hippocampi of *NL-G-F* and *NL-G-F;Lrp6-val* mice at 7 months. Scale bar = 200 µm. D) No differences in plaque number or Aβ coverage in the hippocampus were observed. *NL-G-F* N = 8, *NL-G-F;Lrp6-val* N = 6. Unpaired T-test. E) Quantification of soluble and insoluble Aβ42 by ELISA showed no differences. *NL-G-F* N = 4, *NL-G-F;Lrp6-val* N = 4. Unpaired T-test. Data are represented as mean $\pm$ SEM.

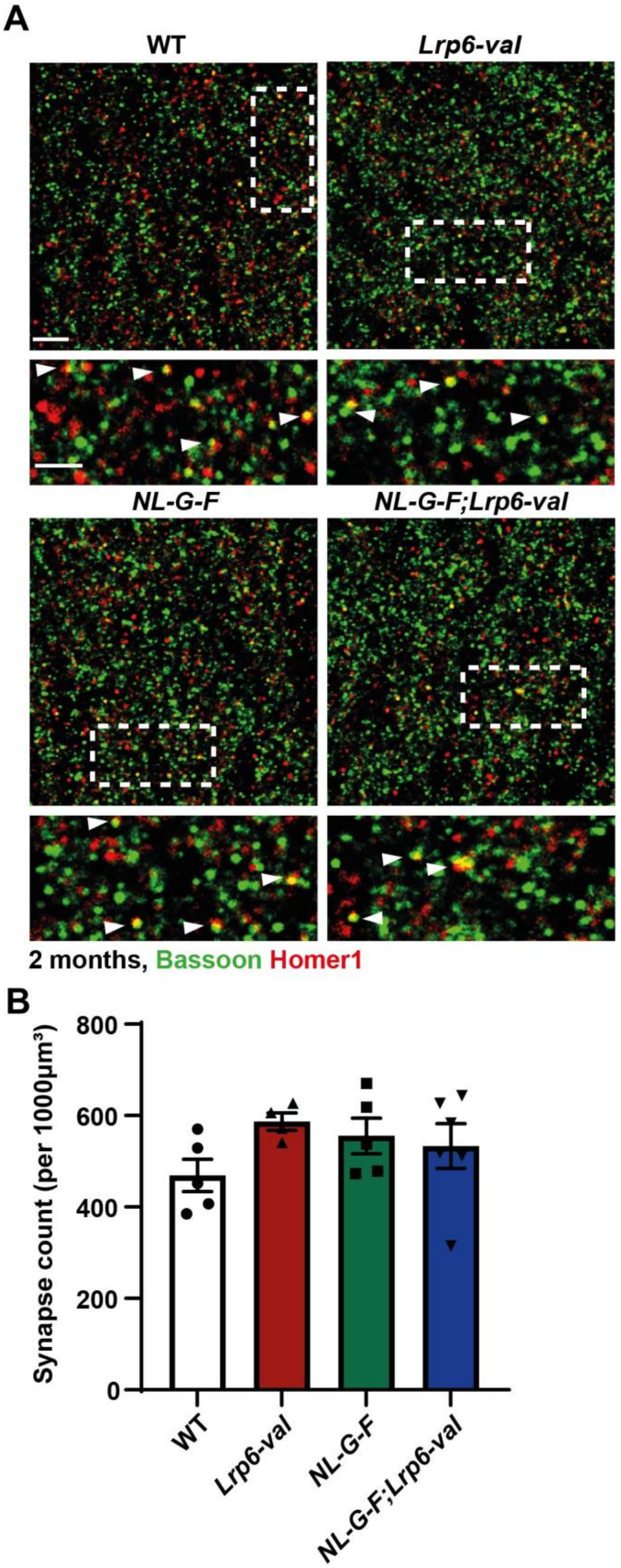
A
WT
Lrp6-val
NL-G-F
NL-G-F;Lrp6-val
2 months, Bassoon Homer1
B
Synapse count (per 1000μm³)
800
600
400
200
0
WT
Lrp6-val
NL-G-F
NL-G-F;Lrp6-val

Figure S7

**Figure S7. *Lrp6-val* does not affect synapse number in *NL-G-F* mice at 2 months.** A) Confocal images of the CA1 SR of WT, *Lrp6-val*, *NL-G-F* and *NL-G-F;Lrp6-val* mice at 2 months showing Bassoon (green) and Homer1 (red) puncta. Scale bar = 3.8 µm. Insets show higher magnification images of synapses. Scale bar = 2 µm B) Quantification of Bassoon and Homer1 puncta and synapse number (co-localised puncta). No differences are detected between any the different genotypes. WT N = 5, *Lrp6-val* N = 4, *NL-G-F* N = 5, *NL-G-F; Lrp6-val* N = 6. One-way-ANOVA with Tukey's post hoc test. Data are represented as mean ± SEM.

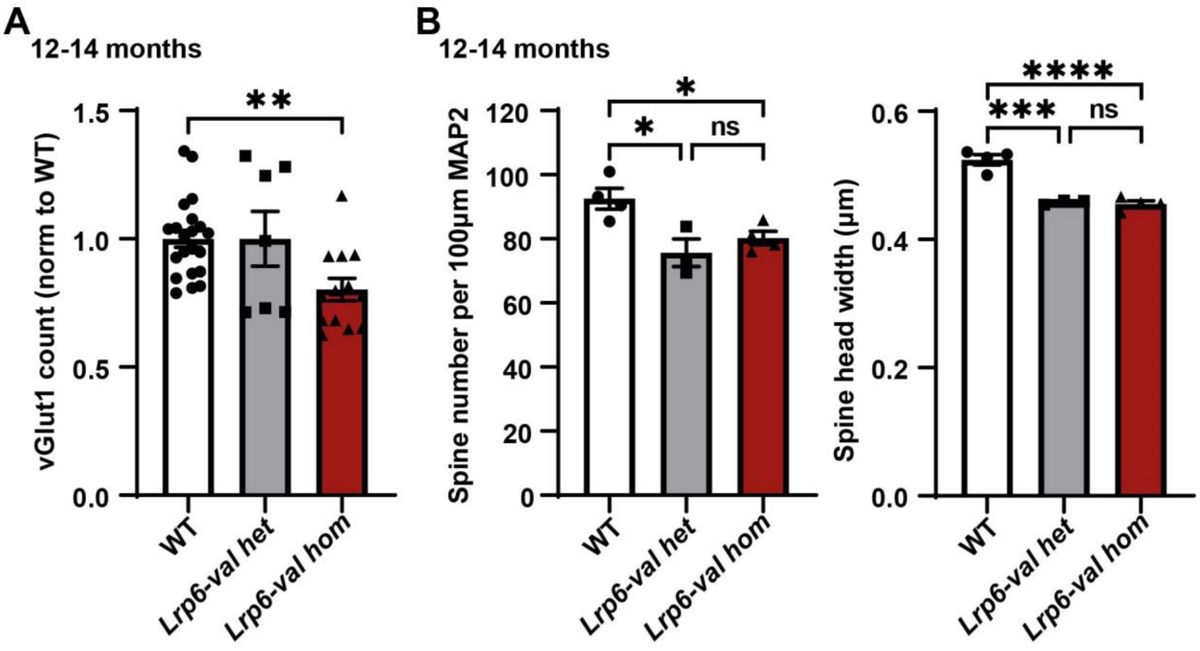


### Figure S8

**Figure S8. Heterozygous *Lrp6-val* mice display post-synaptic defects at 12-14 months.** A) Quantification of vGlut1 puncta in the CA1 stratum radiatum of WT, *Lrp6-val* het and *Lrp6-val* hom mice at 12-14 months. *LRP6-Val* hom mice exhibited fewer vGlut1 puncta. WT N = 10, *Lrp6-val* het N = 4, *Lrp6-val* hom N = 10, 1-3 slices per brain. One-way-ANOVA with Tukey's post hoc test. B) Analyses of dendritic spines in *Lrp6-val* mice crossed to a *Thy1-GFP* reporter line at 12-14 months. Spine number and head width were decreased in both *Lrp6-val* het and *Lrp6-val* hom mice when compared to WT mice. WT N = 4, *Lrp6-val* het N = 3, *Lrp6-val* hom N = 4. One-way-ANOVA with Tukey's post hoc test. Data are represented as mean $\pm$ SEM.
